# Pre-existing chromatin accessibility and gene expression differences among naïve CD4^+^ T cells influence effector potential

**DOI:** 10.1101/2021.04.21.440846

**Authors:** Dakota Rogers, Aditi Sood, HanChen Wang, Jasper J. P. van Beek, Thomas J. Rademaker, Patricio Artusa, Caitlin Schneider, Connie Shen, Dylan C. Wong, Marie-Ève Lebel, Stephanie A. Condotta, Martin J. Richer, Andrew J. Martins, John S. Tsang, Luis Barreiro, Paul Francois, David Langlais, Heather J. Melichar, Johannes Textor, Judith N. Mandl

**Affiliations:** Department of Physiology, McGill University, Montreal, Canada; McGill Research Centre for Complex Traits, McGill University, Montreal, Canada; Immunology-Oncology Unit, Maisonneuve-Rosemont Hospital Research Center, Montreal, Canada; Department of Microbiology, Immunology, and Infectious Disease, Université de Montréal, Montréal, Canada; Department of Medicine, Université de Montréal, Montréal, Canada; Department of Tumor Immunology, Radboud Institute for Molecular Life Sciences, Nijmegen, The Netherlands; Department of Physics, McGill University, Montreal, Canada; Department of Microbiology and Immunology, McGill University, Montreal, Canada; Systems Genomics and Bioinformatics Unit, Laboratory of Immune System Biology, National Institute of Allergy and Infectious Diseases, National Institutes of Health, Bethesda, USA; Department of Genetic Medicine, University of Chicago, Chicago, USA; Department of Human Genetics, McGill University, Montreal, Canada; McGill University Genome Centre, Montreal, QC, Canada

**Keywords:** Cell heterogeneity, CD4^+^ T cells, CD5, T cell differentiation, follicular helper T cells, chromatin landscapes, T cell response, thymus, T cell development

## Abstract

CD4^+^ T cells have a remarkable potential to differentiate into diverse effector lineages following activation. Here, we probed the heterogeneity present among naïve CD4^+^ T cells before encountering their cognate antigen to ask whether their effector potential is modulated by pre-existing transcriptional and epigenetic differences. Using single-cell RNA sequencing, we showed that key drivers of variability are genes involved in T cell receptor (TCR) signaling. Using CD5 expression as a read-out of the strength of tonic TCR interactions with self-peptide MHC, and sorting on the ends of this self-reactivity spectrum, we find that pre-existing transcriptional differences among naïve CD4^+^ T cells impact follicular helper cell (T_FH_) versus non-T_FH_ effector lineage choice. Moreover, our data implicate TCR signal strength during thymic development in establishing differences in naïve CD4 T cell chromatin landscapes that ultimately shape their effector potential.

## Introduction

Heterogeneity is a fundamental property of cellular systems (Altschuler and Wu, 2010; Mayer et al., 2016). Even clonally-derived cell populations exhibit variations in gene expression which impact cell fate decisions (Carter and Zhao, 2020; Chang et al., 2008). Recent single-cell studies examining the epigenome and transcriptome of immune cells have begun to reveal the diversity present among populations thought to be homogeneous and have emphasized the importance of this diversity in the immune response (Brown et al., 2019; Villani et al., 2017; Xie et al., 2020). Such heterogeneity is perhaps nowhere as intimately tied to cellular function as it is in cells of the adaptive immune system. T cell populations comprise a breadth of T cell receptors (TCRs), with individual cells expressing unique TCRs generated by somatic recombination of germline encoded gene segments (Schatz and Ji, 2011). CD4^+^ T cells, in particular, possess a remarkable capacity for diversification into distinct effector lineages following recognition of cognate antigen that are defined by the cytokines they make and the immune cells they act on. Indeed, CD4^+^ T cell effector fate is critical to orchestrating an immune response tailored to the specific pathogen encountered (Zhou et al., 2009; Zhu et al., 2010).

CD4^+^ T cell differentiation relies on dynamic epigenetic, metabolic, and transciptional changes (Almeida et al., 2016; Rodriguez et al., 2015). An early CD4^+^ T cell fate decision is between effector cell subsets (including T helper 1 (T_H_1), T_H_2, T_H_9, and T_H_17) and follicular helper T cells (T_FH_), the latter of which provide essential help for germinal center (GC) formation as well as the affinity maturation of memory B and long-lived plasma cells. This lineage bifurcation into T_FH_ is a multistep process that requires TCR engagement; expression of the lineage-defining transcription factor Bcl6 and concomitant inhibition of Blimp1; and upregulation of surface proteins such as PD-1, CXCR5, and ICOS, enabling migration to and interaction with B cells at the GC border (Crotty, 2019; Qi, 2016; Ruterbusch et al., 2020). Additionally, T_FH_ cells were recently shown to derive from IL-2-producing cells, with IL-2 acting in a paracrine fashion to reinforce non-T_FH_ effector differentiation on responder CD4^+^ T cells (DiToro et al., 2018). Remarkably, a single CD4^+^ T cell clone can expand into a population of multiple helper cell programs after cognate antigen recognition, diversifying into both T_FH_ and non-T_FH_ effector lineages (Becattini et al., 2015; Tubo et al., 2013). CD4^+^ T cell clones are not equal, however. The propensity of a single CD4^+^ T cell to differentiate into one helper subset over another varies between cells expressing distinct TCRs (Cho et al., 2017; Tubo et al., 2013). Whether this clonal variablity in cell fate post-activation is a property determined entirely by TCR signal strength upon cognate antigen encounter, or whether naïve T cells are pre-wired in ways that impact their responses to antigen remains incompletely understood.

It is increasingly appreciated that naïve CD4^+^ T cells already differ prior to antigen stimulation. Despite their actively maintained quiescent state, naïve CD4^+^ T cells remain ready to rapidly respond to antigen (Chapman et al., 2020; Stefanová et al., 2002; Wolf et al., 2020). This is mediated at least in part by tonic TCR signaling through sub-threshold interactions with self-peptide presented on MHC (self-pMHC) that provide survival signals to naïve T cells and keep them poised for activation (Vrisekoop et al., 2014). Several markers have been identified that provide read-outs of the self-pMHC signal strength obtained by naïve T cells. One of these, Ly6C, distinguishes naïve CD4^+^ T cells with high (Ly6C^−^) from low (Ly6C^+^) self-pMHC reactivity (Guichard et al., 2017; Martin et al., 2013). The expression levels of two others, CD5, a surface glycoprotein and negative regulator of TCR signaling, and the orphan nuclear receptor Nur77 (*Nr4a1*), both positively correlate with sub-threshold TCR signal strength (Azzam et al., 2001; Azzam et al., 1998; Moran et al., 2011). Measures of the distribution of CD5 and Nur77 on naïve T cells have revealed that T cell populations span a spectrum of self-reactivities, whereby the upper and lower bounds of self-pMHC reactivity are likely set during thymic development by positive and negative selection (Fulton et al., 2015; Mandl et al., 2013). A combination of all three markers indicates that even within a monoclonal TCR transgenic population, cells vary with regard to basal TCR signal strength (Zinzow-Kramer et al., 2019). Importantly, not only does self-pMHC reactivity impact competition between T cells for homeostatic signals (Vrisekoop et al., 2017), it has also been shown to influence their responses to antigen in specific ways. CD5^hi^ naïve CD4^+^ T cells express higher basal levels of NF*κ*B, phosphorylated TCR-*ζ*, and ERK, make more IL-2 post-activation, preferentially differentiate into regulatory T cells (Tregs), predominate in acute infections, and contribute disproportionately to the memory T cell pool (Fulton et al., 2015; Henderson et al., 2015; Mandl et al., 2013; Matson et al., 2020; Persaud et al., 2014). In contrast, CD5^lo^ naïve CD4^+^ T cells were recently shown to produce more IFN-*γ* upon activation (Sood et al., 2019). Similarly, Ly6C^−^ cells preferentially differentiate into Tregs and T_H_17 cells (Martin et al., 2013). Thus, recent evidence has implicated self-pMHC reactivity of naïve T cells in specific effector response outcomes. However, while these studies examined peripheral T cell function based on self-reactivity, it remains unclear whether T cells are pre-programmed entirely as a result of tonic peripheral TCR signals or whether early encounters with subthreshold ligands in the thymus play a role.

Here, we adopted an unbiased systems approach, combining single-cell (sc) RNA-seq with bulk RNA sequencing (RNA-seq) and an assay for transposase-accessible chromatin using sequencing (ATAC-seq) (Buenrostro et al., 2013), to comprehensively investigate the drivers of heterogeneity among naïve CD4^+^ T cells. We report that individual cell-level biases in the expression of modulators of TCR signal strength, and in T_FH_ versus non-T_FH_ effector lineage choice, are driven, at least in part, by pre-existing transcriptional and epigenetic differences between cells with high (CD5^hi^) versus low (CD5^lo^) self-pMHC reactivity. Unexpectedly, our data reveal that many of the differences in chromatin-accessibility and gene expression among naïve CD4^+^ T cells do not require continuous signaling through the TCR via self-pMHC interactions but are likely a result of variable signal strengths obtained in the thymus during development.

## Results

### TCR signaling induced gene expression differences are drivers of variability among individual naïve CD4^+^ T cells

To define transcriptional differences among naïve CD4^+^ T cells with single cell resolution, we first performed scRNA-seq using CEL-Seq2 (Hashimshony et al., 2016) on a population of 1152 naïve CD4^+^ T cells (sorted from the spleen), of which 697 cells passed quality control tests (**Table S1**). During the cell sorts, we included measures of CD5 protein level expression for each individual T cell given prior studies implicating CD5 as a key read-out of diversity among naïve T cells (Bartleson et al., 2020; Fulton et al., 2015; Henderson et al., 2015; Mandl et al., 2013; Matson et al., 2020; Persaud et al., 2014; Vrisekoop et al., 2017). We verified that among cells in which *Cd5* was detected, transcript counts correlated with measured CD5 protein levels, and drop-outs were more frequent in cells with lower CD5 mean fluorescent intensity (**Figure S1**), consistent with prior evidence for regulation of CD5 at the transcriptional level (Arman et al., 2004; Tung et al., 2001). In total we detected 14,040 genes in our dataset, with an average of 1,389 genes amplified per individual T cell. To investigate genes driving naïve CD4^+^ T cell heterogeneity, we next determined the top 5% most variable genes (total of 716), after accounting for technical variation, frequent non-detection drop-outs, and removing mitochondrial genes (**Table S2**) (Lun et al., 2016). Paralleling a recent study (ElTanbouly et al., 2020), we detected genes known to be important for T cell trafficking between blood and secondary lymphoid organs, and in localization within lymphoid organs including *Cd69*, *S1pr1*, *Sell*, *Klf2*, *Itag4*, *Tln1*, and *Foxo1*, as well as genes involved in TCR signaling, such as *Ptpn6*, *Folr4*, *Il7r*, *Cd4*, *Jun*, *Il2rg*, *Thy1*, *Lck*, *Klf6*, *Bcl2*, *Cd5* and *Nr4a1* (Nur77) (Figure 1A). Interestingly, we also identified a number of genes involved in chromatin modification, including *Dnmt1*, *Hdac4*, *Sirt1*, and *Smc4*, suggestive of possible epigenetic heterogeneity (Figure 1A). Gene ontology (GO) enrichment analyses of the most variable genes identified (Bindea et al., 2009), showed an enrichment of GO terms associated with TCR signaling, including *αβ* T cell activation, regulation of the TCR signaling pathway, regulation of T cell activation, and T cell selection (Figure 1B). We performed a dimensionality reduction with unsupervised uniform manifold approximation (UMAP) analysis based on expression of 55 genes in the GO terms involved in T cell activation from Figure 1B. Overlaying individual cell CD5 protein expression levels showed that the transcriptional state of CD5^lo^ naïve CD4^+^ T cells differed from that of CD5^lo^ naïve CD4^+^ T cells, albeit with considerable overlap among individual cells between the two populations (Figure 1D).

**Figure 1.**
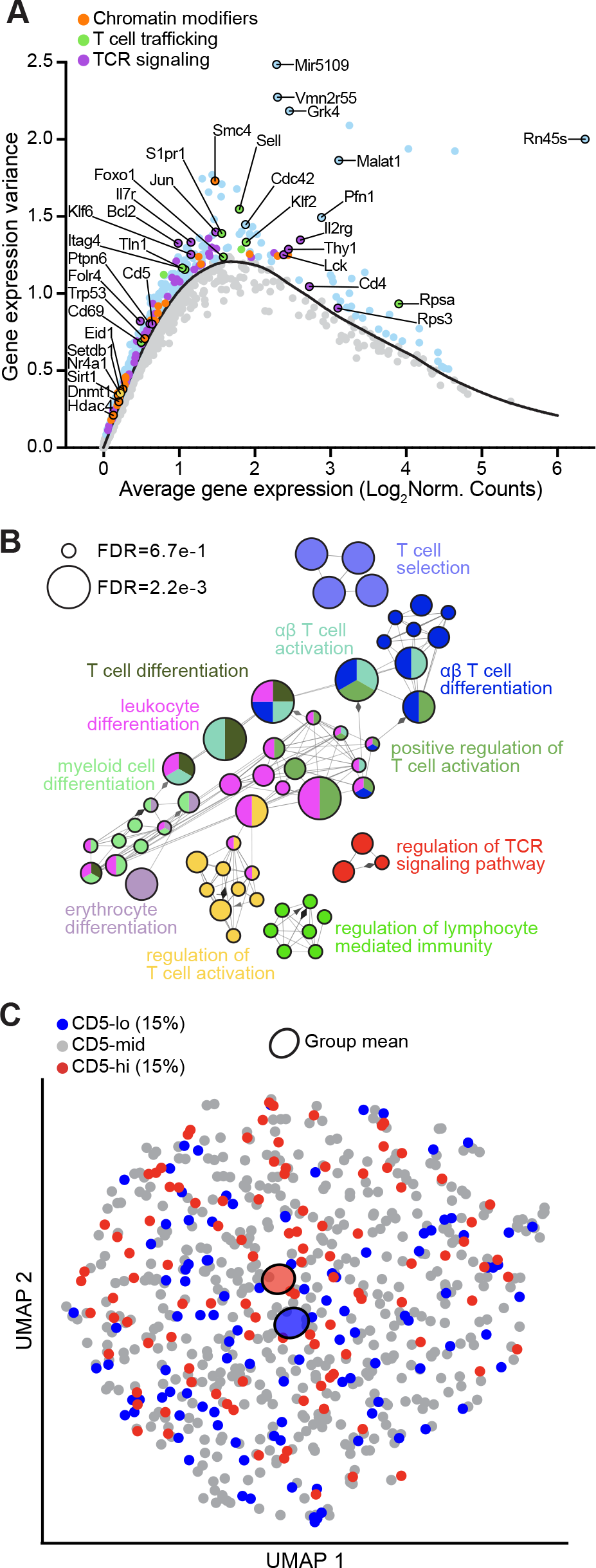
Cellular variability among naïve CD4^+^ T cells at the single cell level is driven by T cell receptor signaling gene expression. **(A)** Plot of the top 5% most variable genes (blue) across individual naïve CD4^+^ T cells (sorted on TCR*β*^+^, CD4^+^, CD8^-^, CD25^-^, CD44^lo^, CD62L^hi^). Select genes involved in TCR signaling (purple), T cell secondary lymphoid organ trafficking (green), and chromatin modification (orange) are labelled. Units for the x- and y-axis are log normalized counts. **(B)** Immune system process gene ontology (GO) enrichment analyses of the top 5% variable genes in naïve CD4^+^ T cells. Circles correspond to unique GO ontology groups with related groups coded in the same color. Circle size reflects enrichment significance (FDR cut-offs as shown). **(C)** UMAP projection of scRNA-seq profiles based on genes that were in the top 5% most variable genes and were also in GO terms involved in T cell activation from B. The surface protein expression of CD5 is overlaid with the 15% CD5^lo^ (blue) and 15% CD5^hi^ (red) naïve CD4 T cells. Shaded colored circles represent CD5^lo^ and CD5^hi^ group means. Each data point represents a single cell. See also Figure S1.

Overall, our data highlight heterogeneity within the naïve CD4^+^ T cell population that is detectable at the transcript-level among single cells and we identified expression differences in genes involved in TCR signaling as drivers of this cellular variability.

### CD5 expression reveals the existence of distinct gene-expression profiles and chromatin landscapes in naïve CD4^+^ T cells

To characterize the transcriptional heterogeneity of naïve CD4^+^ T cells in greater depth, we performed bulk RNA sequencing (RNA-seq), focusing on the ends (top and bottom 15%) of the self-reactivity spectrum as defined by CD5 expression. We used FoxP3^GFP+^ reporter mice as donors for the RNA-seq cell sorts, excluding Tregs from naïve CD4^+^ T cells and including them as an ‘outgroup’ in our initial RNA-seq analyses (Figure 2A, gating strategy and sort purity shown in **Figures S2A** and **S2B**), as CD5 expression is high among Tregs (Ordoñez-Rueda et al., 2009). Given the detection of chromatin modifiers in our scRNA-seq dataset (Figure 1A), we also performed ATAC-seq to investigate the open-chromatin landscape of CD5^lo^ and CD5^hi^ naïve CD4^+^ T cells. After sorting, the difference in surface CD5 expression between CD5^lo^ and CD5^hi^ populations was 5.8 fold, with sorted FoxP3^GFP+^ Tregs having CD5 surface levels 1.7 times lower than the CD5^hi^ cells (Figure 2B).

**Figure 2.**
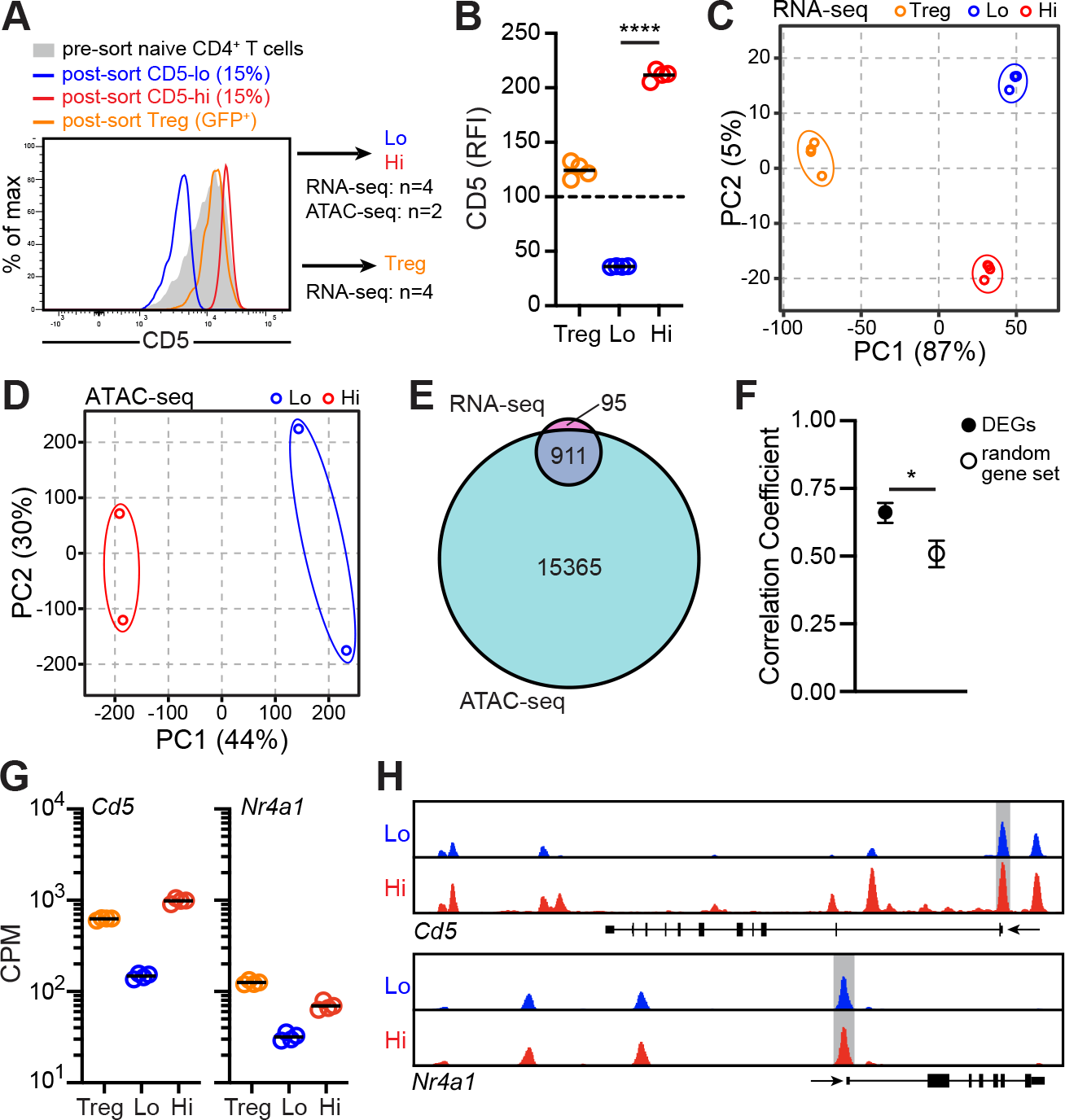
Sorting on CD5 expression reveals that naïve CD4^+^ T cells encompass distinct gene-expression profiles and chromatin landscapes. (**A**) Sorted CD5^lo^, CD5^hi^ naïve CD4^+^ T cells, and regulatory T cell (Treg) populations for RNA- and ATAC-seq. (**B**) Relative fluorescence intensity (RFI) of surface CD5 expression on sorted populations from A, relative to pre-sorted polyclonal naïve CD4^+^ T cells. Lines represent group means. (**C,D**) Principal component analysis (PCA) of populations in A, for RNA-seq (C), or ATAC-seq (D). Numbers in parentheses indicate the variation described by each principal component. (**E**) Venn diagram illustrating overlap in differentially expressed genes (DEG, FDR*≤*0.01) identified by RNA-seq and differentially accessible regions (DARs) identified by ATAC-seq in CD5^lo^ versus CD5^hi^ naïve CD4^+^ T cells. The number of DEG or DAR in each region are indicated. (**F**) Correlation between CD5^hi^ versus CD5^lo^ fold change differences in identified DEGs and corresponding DARs as compared to the correlation from RNA-seq and ATAC-seq within a random gene set. Error bars represent 95% confidence intervals. (**G**) *Cd5* and *Nr4a1* mRNA expression assessed by RNA-seq in populations from A. All group comparisons are significant at FDR<0.01. Lines represent group means. (**H**) ATAC-seq signal profiles across *Cd5* and *Nr4a1* gene loci from one of 2 independent experiments. Promoter regions are highlighted in grey. Statistics: Significantly DEGs (1006) had FDR<0.01 (E), correlation coefficient with *95% CI (F), CPM between CD5^lo^ and CD5^hi^ FDR<0.01 (G). See also Figure S2.

Principal component analyses (PCA) highlighted that at the transcriptional level, CD5^lo^ and CD5^hi^ naïve CD4^+^ T cells clustered into distinct populations (Figure 2C). In addition to being distinguishable from each other in PC2, both the CD5^lo^ and CD5^hi^ populations separated from Tregs in PC1 with respect to their transcriptional programs (Figure 2C). The segregation of CD5^lo^ and CD5^hi^ naïve CD4^+^ T cells was also reflected by their chromatin accessibility profiles, whereby PC1 separated the CD5^lo^ from CD5^hi^ population and PC2 reflected variability between biological replicates (Figure 2D). Overall, CD5^lo^ and CD5^hi^ cells had comparable numbers of accessible peaks and proportions of peaks within exonic, intergenic and intronic regions of the genome (**Figure S2C and S2D**). Importantly, we identified a total of 1,006 differentially expressed genes (DEGs) using an FDR cutoff of < 0.01 between CD5^lo^ and CD5^hi^ naïve CD4^+^ T cells, of which 90% were also found among detected differentially accessible regions (DARs) (Figure 2E). Indeed, there was significantly greater correspondence between the level of gene expression and of open chromatin peak height among DEGs than for a random set of genes (Figure 2F and **S2E**).

Among both the DEGs and DARs identified was CD5 itself, which had a ∼6.7 fold greater transcript expression in CD5^hi^ cells, in line with protein surface expression levels being regulated at the transcriptional level (Fulton et al., 2015) (Figure 2G). In addition, we showed that the *Cd5* locus had a striking reduction in accessible peak number and height in CD5^lo^ compared to CD5^hi^ cells (Figure 2H). It had previously been shown that both transcript and protein levels of Nur77, an immediate-early response gene after TCR stimulation, also reflect the strength of self-pMHC TCR signals obtained by naïve CD8^+^ and CD4^+^ T cells (Fulton et al., 2015; Guichard et al., 2017; Moran et al., 2011). Consistent with this, *Nr4a1* was also among the DEGs (Figure 2G) and, as previously described (Moran et al., 2011), we found that *Nr4a1* expression was greater in Tregs than in CD5^hi^ CD4^+^ T cells. Similar to *Cd5*, the *Nr4a1* locus was more open in CD5^hi^ cells (Figure 2H). Together, our findings reveal considerable differences both at the transcriptional and the chromatin level between CD5^lo^ and CD5^hi^ naïve CD4^+^ T cells.

### Differences in expression of transcriptional regulators and chromatin modifiers may contribute to functional differences among CD5^lo^ and CD5^hi^ naïve CD4^+^ T cells

We next further examined the DEGs identified through bulk RNA-seq and pairwise comparison of the sorted CD5^lo^ and CD5^hi^ naïve CD4^+^ T cell populations. A greater number of the total 1,006 DEGs were upregulated in CD5^hi^ cells (643 genes) than CD5^lo^ cells (363 genes), and, among CD5^hi^ cells, more transcripts were expressed at a fold change of 2 or more compared to CD5^lo^ cells (Figure 3A, **Table S3**). Importantly, despite the high transcript drop-out rates within the scRNA-seq dataset, we nonetheless found strong concordance between the bulk and scRNA-seq datasets with regard to specific genes. For instance, *Cd5*, *Folr4*, *Cd6*, *Nr4a1, Tox*, *Ptpn6,* and *Tcf25* were among genes detected to be more highly expressed in individual CD5^hi^ cells, while *Ly6c1* and *Dntt* were detected at greater levels among individual CD5^lo^ cells (Figure 3B). Indeed, as was also described in a comparable CD5-sorted naïve CD8^+^ T cell dataset (Fulton et al., 2015) and in human CD5^lo^ naïve CD4 T cells (Sood et al., 2021), one of the most differentially expressed transcripts between CD5^lo^ and CD5^hi^ CD4^+^ T cells in our bulk RNA-seq was *Dntt* (encoding the DNA polymerase terminal deoxynucleotidyl transferase, TdT, which inserts non-templated nucleotides during V(D)J TCR rearrangement), at a 14x greater abundance in the CD5^lo^ CD4^+^ T cell population.

**Figure 3.**
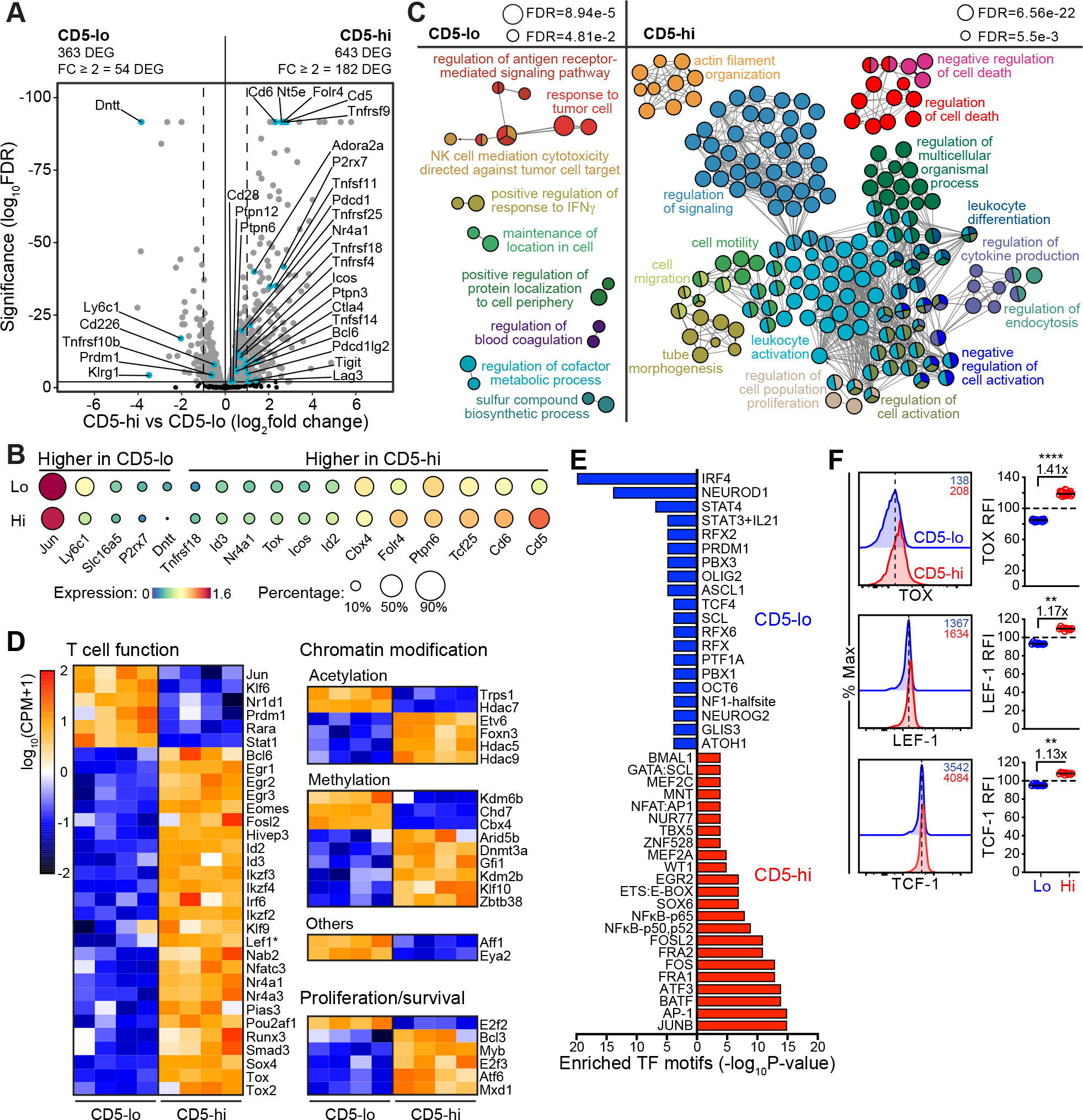
Transcriptional diversity among naïve CD4^+^ T cells suggest functional differences through transcriptional regulators activity and chromatin modifiers. (**A**) Volcano plot of DEGs identified by RNA-seq comparison of CD5^lo^ and CD5^hi^ naïve CD4^+^ T cells. Negative and positive FC values indicate increased expression in CD5^lo^ and CD5^hi^ naïve CD4^+^ T cells, respectively. Significant DEGs (FDR<0.01) are indicated in grey and a subset of DEGs are labeled and highlighted in blue. Total DEG number and DEG with FC ≥ 2 are indicated. Dotted lines are drawn at FC = 2. (**B**) scRNA-seq gene expression level (indicated by colour) and frequency of cells with detectable expression (indicated by circle size) of select DEGs identified from bulk RNA-seq. (**C**) GO enrichment analysis for genes that are upregulated in CD5^lo^ (left) or CD5^hi^ (right) naïve CD4^+^ T cells. Circles correspond to unique GO ontology groups with related groups coded in the same color. Circle size reflects enrichment significance (FDR cut-offs as shown). (**D**) Heatmap of all differentially expressed transcriptional regulators (TR) between CD5^lo^ and CD5^hi^ naïve CD4^+^ T cells, grouped by function. (**E**) Significant (*P*-value <10^-4^) unique enriched TF motifs in DARs from ATAC-seq for CD5^lo^ (blue) and CD5^hi^ (red) naïve CD4^+^ T cells. (**F**) Protein expression of the TRs TOX, LEF-1, and TCF-1 in CD5^lo^ and CD5^hi^ naïve CD4^+^ T cell (RFI is relative to total naïve CD4^+^ T cells). Representative flow cytometry histograms are shown, and data are summarized from 2-5 independent experiments. Dotted lines in histograms denote CD5^lo^ modes; data points in graphs represent individual mice; lines denote group means and average fold differences are indicated for each TR. Statistics: All TRs had FDR<0.01 except *Lef1* (FDR=0.012) (C), Wilcoxon matched-pairs signed rank test (E). \*\**p*<0.01, \*\*\*\**p*<0.0001. See also Figure S3.

Given the greater *Dntt* expression in CD5^lo^ cells among both naïve CD4^+^ and CD8^+^ T cell subsets, we asked whether other DEGs we identified from the CD5^lo^ versus CD5^hi^ comparison within CD4^+^ T cells were shared with naïve CD8^+^ T cells sorted on CD5 expression levels. Interestingly, in line with the narrower CD5 distribution among CD8^+^ T cells (Mandl et al., 2013), far fewer DEGs with a fold change cut-off ≥2 were previously identified among CD5^lo^ versus CD5^hi^ CD8^+^ T cells (total of 57) than in our CD4^+^ dataset (total of 236 DEGs). The overlap between CD5^lo^ versus CD5^hi^ DEGs in CD4^+^ T cells compared to CD8^+^ T cells was only 10%, and there were 33 CD5hi/lo CD8^+^ DEGs that were not detected or not significantly differently expressed among naïve CD4^+^ T cells including *Xcl1*, *Cxcr3*, *Ptpn4*, and *Tbx21* (**Figure S3A**). The majority of the 24 DEGs shared between CD4^+^ and CD8^+^ T cells showed concordance such that transcripts with greater expression in CD5^hi^ CD8^+^ cells were also upregulated in CD5^hi^ CD4^+^ T cells relative to CD5^lo^ cells, including *Itih5*, *Eomes*, *Cd200*, and *Ikzf2* (**Figure S3B**). However, two genes, *Ly6c1* and *Ddc,* did not follow this trend and expression differences were opposite when comparing naive CD8^+^ and CD4^+^ T cells (**Figure S3B**). Also, unlike CD5^hi^ CD8^+^ T cells, CD5^hi^ CD4^+^ T cells were not larger in size and did not express greater surface CD44 levels (**Figure S3C**). Thus, overall, there were few parallels between naïve CD4^+^ T cells and CD8^+^ T cells sorted on CD5 expression with regard to the specific DEGs identified.

To investigate patterns in DEGs increased among CD5^lo^ or CD5^hi^ CD4^+^ T cells, we performed GO enrichment analyses. In the CD5^lo^ population we found an enrichment of GO terms associated with tumor-mediated immunity and regulation of IFN-*γ* responses (Figure 3C). The latter is consistent with recent work showing that CD5^lo^ CD4^+^ T cells produce more IFN*γ* than CD5^hi^ cells upon activation (Sood et al., 2019). In contrast, in the CD5^hi^ population, gene networks involved in leukocyte activation, regulation of signaling, and cell migration were strongly enriched (Figure 3C). In line with this, gene set enrichment analysis (GSEA) indicated that genes associated with CD4^+^ T cell activation and effector responses (Gottschalk et al., 2012; Hale et al., 2013), were significantly overrepresented in the CD5^hi^ population, whereas genes associated with CD4^+^ T cell memory were overrepresented in the CD5^lo^ naïve CD4^+^ T cell population (**Figure S3D**). Of note, genes involved in the active maintenance of a quiescent state among naïve T cells, including *Klf2*, *Tgfbr2, Btg1, Btg2, Tob1*, *Foxo1*, *Foxo3*, *Foxp1*, *Slfn2*, *Tsc1*, and *Tsc2* (Chapman et al., 2020; ElTanbouly et al., 2020; Hamilton and Jameson, 2012; Yusuf and Fruman, 2003), were detected in both CD5^lo^ and CD5^hi^ naïve CD4^+^ T cells and were not differentially expressed between the two groups (**Table S3, S4**). Moreover, no cytokines were among the identified DEGs, with the exception of *Il16* which is known to be constitutively expressed in naïve CD4^+^ T cells (Ren et al., 2005) and was slightly increased in CD5^lo^ cells (**Table S3, S4**). Indeed, the chromatin loci for effector cytokines such as *Ifng*, *Il5*, *Il17a*, *IL21*, and *Il10* were closed; only the transcription start site for *Il2* had a slightly greater accessible peak in CD5^hi^ cells (**Figure S3E**). These data indicated that although CD5^hi^ cells were enriched for gene networks associated with activation, both gene expression and accessible chromatin regions were largely consistent with an equally quiescent and non-differentiated cell state among naïve CD4^+^ T cells with different self-reactivities.

Quiescence exit occurs when naïve T cells obtain antigen stimulation and co-stimulation, but before the first cell division (Chapman et al., 2020). Once they receive activating signals, T cells undergo chromatin remodeling, transcriptional changes, and ultimately effector differentiation. To understand whether naïve CD4^+^ T cells were poised to respond differently to activation as a function of the transcriptome and chromatin accessibility differences we had identified, we first investigated whether they differed in expression of transcriptional regulators (TRs) important to T cell activation and function. For this we used a predefined list of 1680 known or putative TRs (Mingueneau et al., 2013). Among DEGs, we detected 31 TRs upregulated in CD5^hi^ cells involved in T cell proliferation or survival (*Atf6*, *Myb*, and *Bcl3*), and T cell activation or differentiation (*Egr1*, *Egr2*, *Egr3*, *Nfatc3*, *Ikfz3*, *Ikzf4*, *Tox*, *Tox2*, *Nr4a1*, *Nr4a3*, *Klf9*, *Lef1*, *Bcl6*, *Eomes*, *Irf6*, *Id2*, and *Id3*) (Figure 3D). We also detected 21 TRs that mediate chromatin modifications such as acetylation (*Hdac5*, *Hdac9*, and *Etv6*) and methylation (*Dmnt3a*, *Klf10*, *Kdm2b*, and *Gfi1*) (Figure 3D), as well as TRs without clearly defined roles in T cells (**Figure S3F**). Twenty TRs were enriched in CD5^lo^ cells including transcriptional repressors (*Hdac7, Nr1d1, Prdm1, Rara,* and *Trps1*) (Figure 3D and **S3F**).

We next asked if there were TR binding motifs enriched among chromatin peaks that were unique to either CD5^lo^ or CD5^hi^ cells. Indeed, CD5^hi^ cells were enriched in binding motifs for transcription factor networks downstream of TCR activation, including AP-1 and JNK transcription factors, FOS, FOSL2, AP-1, JUNB, as well as NUR77, NFAT, and NF*κ*B (Figure 3E and **S3G**). Conversely, CD5^lo^ cells were enriched in binding motifs for IRF-4 and BLIMP-1 (PRDM1), both of which promote non-T_FH_ effector differentiation (Johnston et al., 2009; Krishnamoorthy et al., 2017). Notably, while not represented in TR binding motif analysis, TOX and TOX2 were among identified differentially upregulated TRs in CD5^hi^ cells, which function downstream of the TCR through nuclear factor of activated T-cells (NFAT) signaling (Khan et al., 2019; Scott et al., 2019; Seo et al., 2019) and were recently shown to promote T_FH_ differentiation by increasing Bcl6 expression through *Tcf7* expression and enhanced chromatin accessibility of TCF-1 and LEF-1 bound regions of the *Bcl6* locus (Xu et al., 2019). Thus, we asked whether TOX, LEF-1, and TCF-1 expression was greater at the protein level in CD5^hi^ cells, as suggested by their increased mRNA levels (Figure 3F). While the detected differences were small (1.3-1.4 fold), they were robust across mice, were not observed in unstained controls (**Figure S3H**), and corresponded to increased chromatin accessibility in the loci for *Tox*, *Tox2*, *Lef1*, and *Tcf7* in CD5^hi^ cells (**Figure S3I**).

Together, our data suggest the possibility that a network of TRs and unique epigenetic profiles results in differences in cell states among naïve CD5^lo^ and CD5^hi^ CD4^+^ T cells impacting their function and/or differentiation upon activation, particularly with regard to the early T_FH_ vs. non-T_FH_ bifurcation.

### Pre-existing expression differences in regulators of TCR signaling among naïve CD4^+^ T cells are maintained after activation

To further investigate the cell signaling and lymphocyte activation signatures in CD5^hi^ CD4^+^ T cells, we first curated a list of DEGs involved in the regulation of T cell activation. CD5^hi^ cells had increased expression of genes involved in co-stimulatory pathways, such as *Icos*, *Rankl*, *Itgb2*, and *Gitr* (Figure 4A). More predominantly, CD5^hi^ cells had a higher expression of genes involved in the negative regulation of TCR signaling or T cell activation, including *Cd6*, *Nt5e* (CD73), *Ptpn6* (SHP-1), *Ctla4*, *Pdcd1* (PD-1), *Btla, IL10ra, P2rx7, Nrp1, Nrp2, Cd200,* and *Adora2a* (Figure 4A). Of note, the checkpoint regulator VISTA recently identified in naïve CD4^+^ T cells was not detected in our dataset (ElTanbouly et al., 2020). Given the role of negative regulators in T cell exhaustion during chronic antigen stimulation, and perhaps indicative of a greater frequency and/or strength of self-pMHC signals obtained by CD5^hi^ T cells, GSEA identified an enrichment in exhaustion-associated genes such as *Tox*, *Tox2*, *Tigit*, *Lag3*, *Ctla4*, and *Cblb* among CD5^hi^ cells (**Figure S4A**).

**Figure 4.**
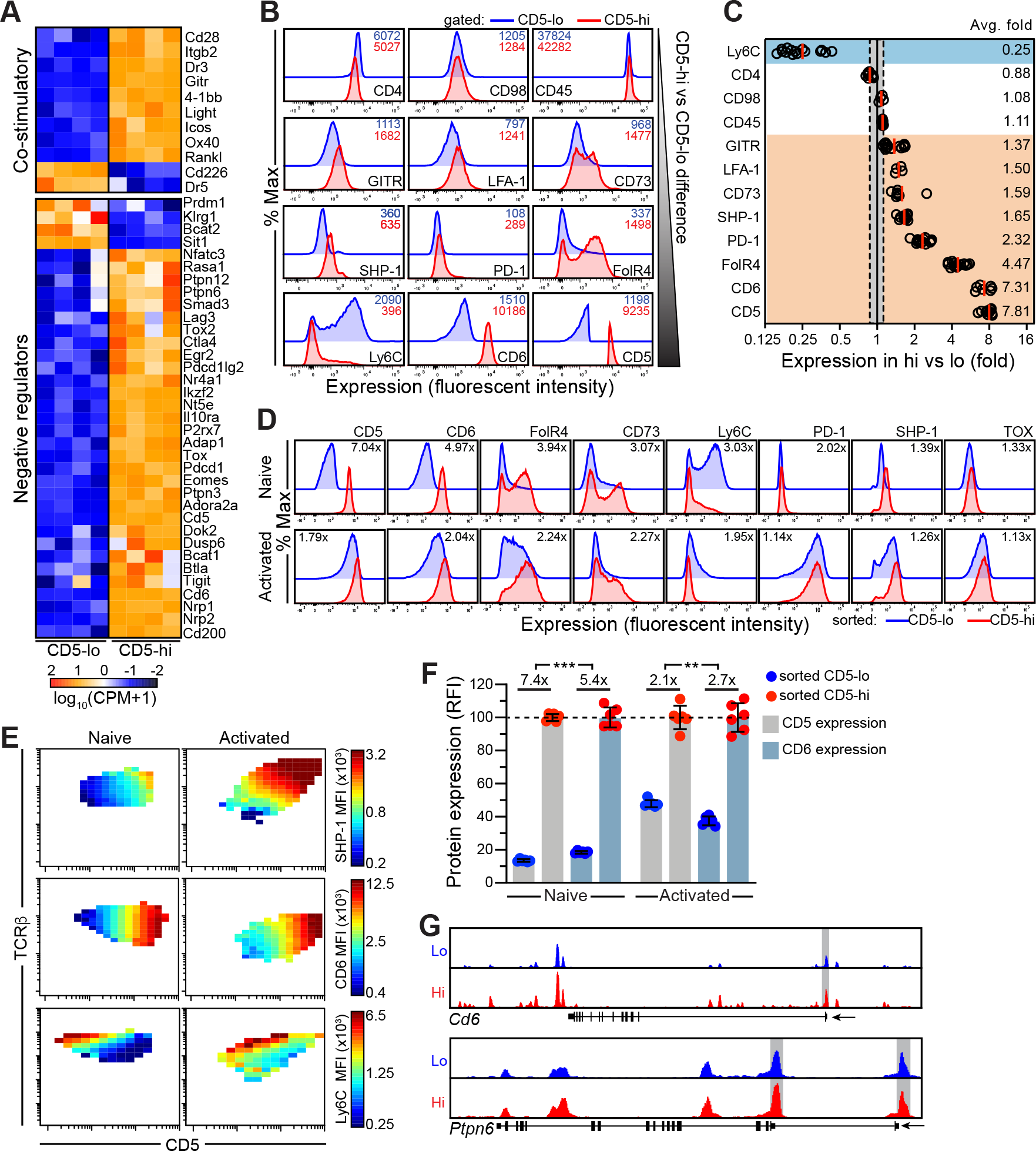
Pre-existing transcriptional and protein differences among naïve CD4^+^ T cells are maintained post-activation. (**A**) Heatmap with unsupervised clustering of curated list of DEGs involved in positive and negative regulation of T cell activation and T cell receptor signaling. (**B**) Representative histograms of protein expression (measured by flow cytometry) of select DEGs identified from bulk RNA-seq analyses comparing CD5^lo^ and CD5^hi^ naïve CD4^+^ T cells. Numbers in top right of histograms indicate fluorescent intensity in CD5^lo^ (blue) and CD5^hi^ (red) populations. (**C**) Summary of fold expression differences of proteins measured in B that were significantly different between CD5^hi^ and CD5^hi^ groups from 2-5 independent experiments. Non-significant DEGs (CD4, CD98, and CD45) were used as controls to establish a cut-off for biologically significant protein fold differences; blue shading indicates greater expression in CD5^lo^ and orange shading indicates greater expression in CD5^hi^. Each data point is from an individual mouse; red lines denote group means. (**D**) Representative histograms of protein expression measured by flow cytometry in sorted 15% CD5^lo^ and CD5^hi^ naïve CD4^+^ T cells pre- and 24 hours post-activation (CD44^hi^CD62L^lo^ or CD44^hi^CD25^+^) with anti-CD3/CD28. Numbers in top of histograms indicate fold difference between CD5^lo^ (blue) and CD5^hi^ (red) populations. (**E**) 3-dimensional single-cell flow cytometry analysis for SHP-1, CD6, and Ly6C expression among naive CD4^+^ T cells pre- and 24 hours post-activation (CD44^hi^CD25^+^). Color scale represents MFI of proteins of interest (SHP-1, CD6, or Ly6C) in bins of at least 10 cells across the full protein expression spectrums of CD5 (x-axis) and TCR*β* (y-axis). (**F**) Naïve CD4^+^ T cells were sorted into CD5^lo^ and CD5^hi^ populations, and CD5 and CD6 protein expression measured pre- and post-activation (CD44^hi^CD25^+^) with anti-CD3/CD28. Representative histograms (top) and data summarized from 2 independent experiments with relative fluorescent intensity normalized to CD5 or CD6 expression in the CD5^hi^ population (bottom). Each data point represents 4-5 pooled mice; error bars represent mean ± S.D. (**G**) ATAC-seq signal profiles across *Ptpn6* and *Cd6* gene loci shown from one of 2 independently sorted CD5^lo^ and CD5^hi^ naïve CD4^+^ T cell samples. Promoter regions are highlighted in grey. Statistics: All genes had FDR<0.01 (A), paired t test (G). \*\**p*<0.01, \*\*\**p*<0.001. See also Figure S4.

Corroborating our gene expression data, we observed expression differences between CD5^lo^ and CD5^hi^ cells also at the protein level for a subset of the regulators of TCR signaling including GITR, LFA-1 (integrin *α*L*β*2), CD6, FolR4, PD-1, and CD73 (varying from 1.4 to 7.8 fold) (Figure 4B and 4C). Interestingly, signaling through LFA-1 was recently shown to promote Bcl6 expression and be required for T_FH_ differentiation following activation (Meli et al., 2016). Notably, some of the other genes modulating the TCR signal have also been shown to be highly expressed by T_FH_ cells and play a role in driving T_FH_ differentiation, including *Tox*, *Tox2*, *FolR4*, and *Icos*, all of which are expressed at greater levels in CD5^hi^ naïve CD4^+^ T cells (Figure 4A). In addition, we confirmed greater protein expression in CD5^hi^ cells of the soluble hematopoietic phosphatase, SHP-1, a negative regulator of TCR signaling that modulates T cell sensitivity and responsiveness to antigen (Feinerman et al., 2008; Stefanová et al., 2003), and associates with CD5 and CD6 (Blaize et al., 2020; Gonçalves et al., 2018), and other negative regulators of TCR signaling such as PD-1, BTLA, and CTLA-4 (Lorenz, 2009) (Figure 4B and 4C). Further, as expected based on published data, CD5^lo^ cells expressed higher levels of Ly6C (Martin et al., 2013). Of note, the non-significant DEGs CD4, CD98 (LAT1), and CD45 were used as controls to establish a cut-off for biologically significant protein fold differences (Figure 4B and 4C). Together, these data indicated that at least for the subset of tested DEGs, transcriptional differences among naïve CD4^+^ cells correlated with differences in protein expression. The increased expression of negative regulators of T cell signaling in CD5^hi^ CD4^+^ T cells in particular may play a role in preventing cells with high self-pMHC reactivity from becoming overtly autoreactive. Indeed many of the negative regulators detected to be increased in CD5^hi^ cells have previously also been shown to be important in tolerance to self-antigens (Kalekar et al., 2016).

Given the importance of levels of TCR signal regulators in the response of T cells to cognate antigen (Feinerman et al., 2008), we next asked whether the differential expression of these regulators among naïve CD4^+^ T cells are maintained following TCR stimulation. We found that polyclonal sorted CD5^lo^ and CD5^hi^ naïve CD4^+^ T cells stimulated with anti-CD3/CD28, which bypasses individual TCR affinity differences for specific agonist peptides, retained expression differences 24 hours after stimulation, albeit for some proteins at reduced fold differences between CD5^lo^ and CD5^hi^ cells (Figure 4D and **S4B**). In unsorted polyclonal CD4^+^ T cells, SHP-1 and CD6 expression increased with greater CD5 levels, and this relationship was maintained following activation (Figure 4E). In contrast, as was shown previously (Martin et al., 2013), Ly6C followed a bimodal distribution, which largely disappeared after activation with most cells becoming Ly6C-negative (Figure 4D **and 4E**). In addition, the transcription factor TOX, which drives expression of exhaustion-associated genes (Scott et al., 2019; Seo et al., 2019), followed a similar expression pattern as SHP-1 across the full spectrum of CD5, but the expression range was reduced post-activation (**Figure S4C**). Mutual information analysis confirmed that CD5 and CD6 were much better predictors of SHP-1 expression than were TCR*β* levels, both pre- and post-activation (**Figure S4D and S4E**), suggesting that self-reactivity likely plays an important role in tuning TCR signal strength during cognate antigen stimulation.

*Cd5* and *Cd6* are gene homologs that likely arose from a duplication event, are both located on the same region on chromosome 11 in humans and 19 in mice, and have functional similarities (Lecomte et al., 1996; Padilla et al., 2000). Our data indicated a tight correlation between CD5 and CD6 levels, with CD5 expression being a robust predictor of CD6 expression both pre- and post-activation (Figure 4E and **S4E**), consistent with previously published data (Richards et al., 2015). Interestingly, the fold change difference in CD6 expression on sorted CD5^hi^ and CD5^lo^ CD4^+^ T cells remained slightly greater post-activation than CD5, suggesting that CD6 might be a useful marker to more reliably infer the self-pMHC reactivity of naïve CD4^+^ T cells after activation (Figure 4F). The maintenance of differences among naïve CD4^+^ T cells in the expression of regulators of TCR signaling even following a strong TCR stimulus raised the possibility that these might be regulated through epigenetic modifications rather than modulated entirely by signal strength through the TCR during homeostatic self-pMHC interactions or stimulation with cognate antigen. Indeed, similarly to our observed differences in chromatin accessibility of the *Cd5* locus (Figure 2H), the *Cd6* and *Ptpn6* loci showed greater peak accessibility in CD5^hi^ cells (Figure 4G). In summary, we corroborated expression differences between CD5^lo^ and CD5^hi^ naïve CD4^+^ T cells at the protein-level for genes important in tuning TCR signal strength and showed that these differences were maintained even after strong TCR stimulation.

### CD5^hi^ cells have a greater propensity to develop into T_FH_ cells than CD5^lo^ naïve CD4^+^ T cells

Given that the DEGs we identified in the CD5^lo^ and CD5^hi^ comparison of naïve CD4^+^ T cells included both TRs and surface molecules known to be involved in T_FH_ differentiation, all of which were expressed more highly in the CD5^hi^ population, we asked whether these pre-existing transcriptional differences might play a role in the early T_FH_ and non-T_FH_ lineage choice following cognate antigen encounter. Indeed, GSEA showed that CD5^hi^ naïve CD4^+^ T cells were significantly enriched for T_FH_ and germinal center (GC) T_FH_ cell gene signatures (Figure 5A). Across all replicates, CD5^hi^ cells expressed higher levels of *Pdcd1*, *Cxcr5*, and *Bcl6*, while CD5^lo^ cells expressed higher levels of the T_FH_ repressor *Prdm1* (Blimp-1) (Figure 5B). Moreover, the *Pdcd1*, *Cxcr5*, and *Bcl6* loci had greater chromatin accessibility in CD5^hi^ naïve CD4^+^ T cells (Figure 5C). These data suggested the possibility of a pre-existing disposition among CD5^hi^ naïve CD4^+^ T cells to become T_FH_ cells relative to their CD5^lo^ counterparts. In support of this hypothesis, CD5^hi^ naïve CD4^+^ T cells were shown to produce more IL-2 than CD5^lo^ cells post-stimulation (Persaud et al., 2014), and recent data has highlighted the importance of early IL-2 production in T_FH_ lineage choice, with IL-2 producers becoming T_FH_ cells and paracrine IL-2 signaling reinforcing non-T_FH_ lineage commitment of CD4^+^ T cells obtaining weaker TCR signals (Ballesteros-Tato et al., 2012; DiToro et al., 2018).

**Figure 5.**
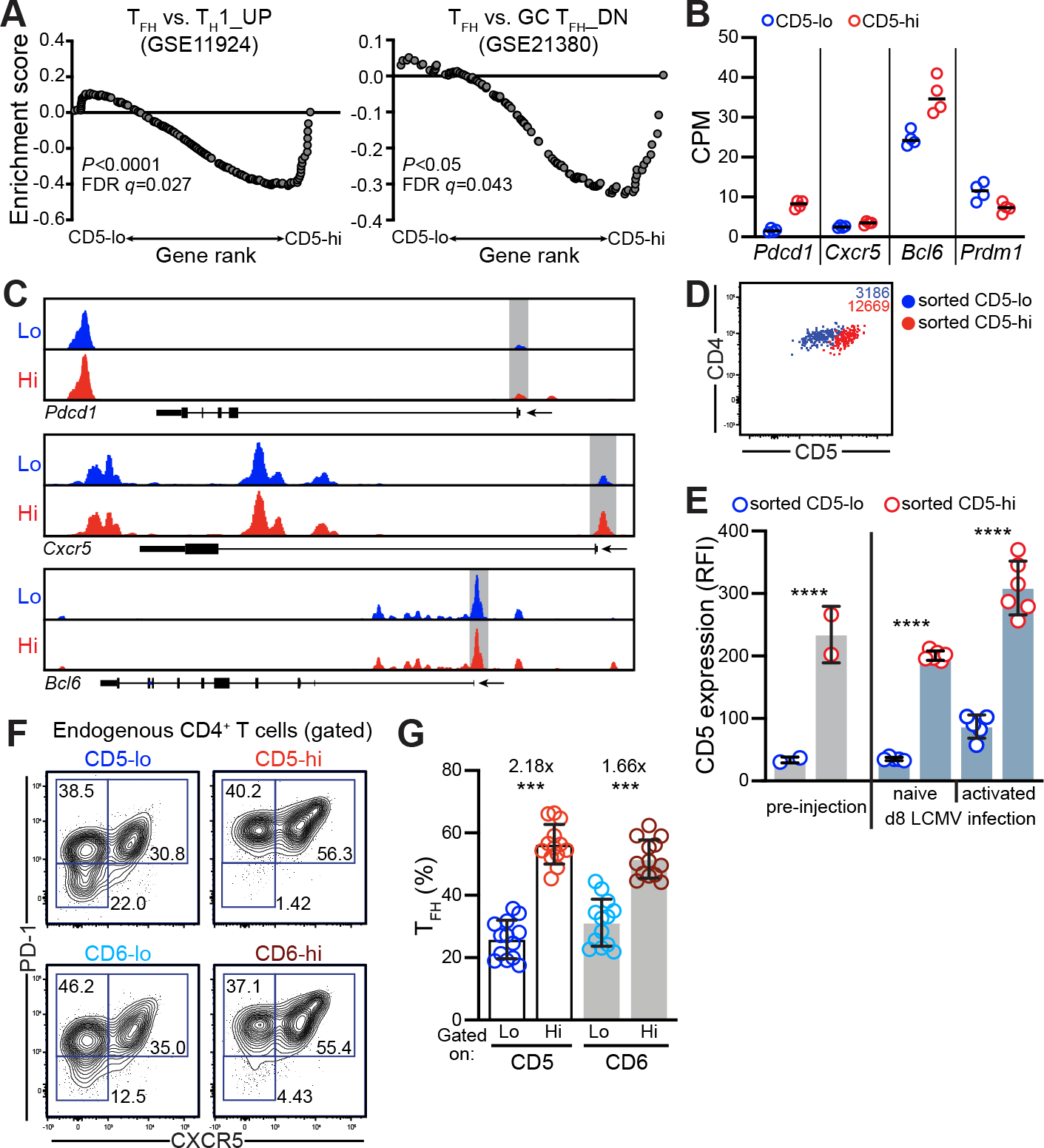
CD5^hi^ naïve CD4^+^ T cells are enriched for T_FH_-associated genes and have a greater T_FH_ differentiation potential upon infection than CD5^lo^ T cells. (**A**) Gene-set enrichment analysis (GSEA) of T**_FH_** signatures enriched in CD5^lo^ and CD5^hi^ naïve CD4^+^ T cells. (**B**) *Pdcd1*, *Cxcr5*, *Bcl6*, and *Prdm1* mRNA expression from bulk RNA-seq for sorted CD5^lo^ and CD5^hi^ naïve CD4^+^ T cell populations. Lines indicate group means. (**C**) ATAC-seq signal profiles across *Pdcd1*, *Cxcr5*, and *Bcl6* gene loci from one of 2 independently sorted CD5^lo^ and CD5^hi^ samples. Promoter regions are highlighted in grey. (**D,E**) Naïve 15% CD5^lo^ and CD5^hi^ CD4^+^ T cells were sorted and adoptively transferred into recipients that were infected 1 day later with LCMV. Representative flow cytometry plot of activated (CD44^hi^) transferred CD5^lo^ and CD5^hi^ CD4^+^ T cells with numbers in top right of dot plots indicating CD5 MFI of sorted CD5^lo^ (blue) and CD5^hi^ (red) populations (D), and summary of CD5 RFI (relative to endogenous total naïve CD4^+^ T cells) of sorted CD5^lo^ and CD5^hi^ naïve or activated CD4^+^ T cells (E). In E, each data point is from an individual recipient mouse; error bars represent mean ± S.D. (**F,G**) Activated (CD44^hi^) CD4^+^ T cells isolated on day 8 post LCMV infection were gated on the top and bottom 15% CD5- or CD6-expressing cells and the percent T_FH_ determined. Representative flow cytometry plots are shown, with numbers indicating percent cells within each gate (F), and data summarized from 2 independent experiments with each data point being from an individual mouse (G). Statistics: All genes had FDR<0.01 except *Cxcr5* (FDR=0.044) (B), one-way ANOVA with Tukey’s multiple comparisons (E), Wilcoxon matched-pairs signed rank test (G). ***p<0.001, ****p<0.0001. See also Figure S5.

To directly assess whether CD5^hi^ naïve CD4^+^ T cells gave rise to a greater proportion of T_FH_ cells *in vivo* than CD5^lo^ cells, we first sorted 15% CD5^lo^ and 15% CD5^hi^ polyclonal naïve CD4^+^ T cells, adoptively transferred 6-10×10^6^ cells of each population into separate recipient groups, and infected recipients with LCMV, which elicits a robust T_FH_ response (Fahey et al., 2011). There are only an estimated 8 LCMV-GP66 specific CD4^+^ T cells per million naïve CD4^+^ T cells (Jenkins and Moon, 2012; Nelson et al., 2015), and even assuming additional antigen-specific CD4^+^ T cells, this would mean that very few antigen specific T cells would be among the transferred sorted populations. Indeed, as expected, the proportion of activated CD4^+^ T cells among the transferred cells 8 days post infection was highly stochastic. However, gating on activated transferred cells showed that, as in our *in vitro* activation assays, the CD5 expression difference between CD5^lo^ and CD5^hi^ CD4^+^ T cells was maintained (Figure 5D and 5E).

As an alternative approach, we asked whether we would be able to better assess differences in T_FH_ differentiation potential by transferring sorted populations into TCR*β*^-/-^ recipients. Given the lack of competitor T cells in these recipients, we first investigated whether this would impact T_FH_ lineage choice in a population of CD4^+^ T cells with a fixed antigen-specific TCR. We performed adoptive transfers of LCMV-specific TCR transgenic (Tg) SMARTA CD4^+^ T cells into both wild-type (WT) and TCR*β*^-/-^ recipients, infected them 1 day later with LCMV and then assessed the response on day 8 post infection (**Figure S5A**). Overall, the clonal expansion of SMARTA TCR Tg cells was greater in the TCR*β*^-/-^ mice (**Figure S5B**). Interestingly, we found that when SMARTA TCR Tg cells were transferred into TCR*β*^-/-^ recipients, CD5 surface expression was significantly increased compared to cells transferred into WT mice, suggesting that SMARTA Tg cells were receiving stronger TCR signals upon cognate antigen encounter in the absence of other competitor T cells, as has been previously shown to be the case in lymphopenia-induced expansion (Vrisekoop et al., 2017) (**Figure S5C**). Importantly, the percent of T_FH_ cells among transferred SMARTA TCR Tg cells was halved in the infected TCR*β*^-/-^ compared to WT recipients (**Figure S5D**). Given the role of IL-2 in T_FH_ differentiation and the modulation of IL-2 production by TCR signal strength (DiToro et al., 2018), we postulated that the decreased T_FH_ differentiation of SMARTA TCR Tg cells in TCR*β*^-/-^ mice was a result of greater IL-2 production mediated by greater TCR signaling. Indeed, we observed that SMARTA TCR Tg cells produced significantly more IL-2 and had greater CD25 (IL-2R*α*) surface expression in the infected TCR*β*^-/-^ mice (**Figure S5E**). Due to the impact of the lack of other T cells in the TCR*β*^-/-^ mice on T_FH_ frequency post infection, our read-out of interest, and the likelihood that the TCR signal strength modulation in TCR*β*^-/-^ mice is not equal between cells of high and low self-reactivity (Vrisekoop et al., 2017), we concluded that this experimental approach would not accurately enable us to identify differences between CD5^lo^ and CD5^hi^ polyclonal naïve CD4^+^ T cells with regard to their T_FH_ differentiation potential. Indeed, a recent study showed that sorted Nur77^hi^ naïve CD4^+^ cells differentiated into T_FH_ cells less than Nur77^lo^ naïve CD4^+^ cells, but utilized TCR*α*^-/-^ mice as recipients for sorted populations and did not account for concomitant changes in IL-2 production and consumption which track with tonic signal strength (Bartleson et al., 2020).

Instead, based on the observation that CD5 expression levels remained detectably different between sorted and transferred CD5^lo^ and CD5^hi^ populations, we quantified T_FH_ differentiation among activated CD4^+^ T cells expressing high or low CD5 levels 8 days post-LCMV infection. We found that there was a ∼2 fold increase in T_FH_ differentiation within the top 15% CD5^hi^ activated CD4^+^ T cell population compared to the bottom 15% CD5^lo^ cells (Figure 5F, **5G**, and **S5F**). Moreover, among CD5^hi^ cells, T_FH_ (PD-1^hi^ CXCR5^+^) and PD-1^hi^ CXCR5^−^ cells expressed greater levels of PD-1 than their CD5^lo^ counterparts (**Figure S5G**). Consistent with CD6 expression levels as another read-out of basal self-pMHC signal strength similar to CD5, we found that T_FH_ cells were also overrepresented among CD6^hi^ CD4^+^ T cells compared to CD6^lo^ cells, with a fold difference similar to CD5^hi^ vs CD5^lo^ (1.66-fold) and had increased PD-1 expression over their CD6^lo^ counterparts (Figure 5F**-G, S5F-G**). In line with an increased T_FH_ population deriving from CD5^hi^ CD4^+^ T cells, CD5^hi^ T_FH_ also expressed higher levels of TOX, LEF-1, and TCF-1 (**Figure S5H**). These transcription factors have been shown to play an essential role in early cell fate decisions and regulate the development of T_FH_ cells by modulating the expression of several T_FH_ associated genes such as *Pdcd1*, *Cxcr5*, *Bcl6*, *Icos*, *Il6ra*, and *Tcf7* (Choi et al., 2015; Xu et al., 2015; Xu et al., 2019). Together, our data suggested that the T_FH_ and non-T_FH_ cell fate decision is altered by pre-existing differences present in the naïve CD4^+^ T cell population prior to foreign antigen encounter.

### Removing naïve CD4^+^ T cells from continuous self-pMHC interactions reveals gene expression differences that are independent of post-thymic self-ligand recognition

Previous work showed that continuous tonic self-pMHC signals obtained by naïve T cells in the periphery facilitate antigen recognition by maintaining partial TCR*ζ*-chain phosphorylation, and through polarization of the TCR and its signaling components (Stefanová et al., 2002). Interrupting signals from self-pMHC interactions for as little as 30 minutes led to a loss of sensitivity to cognate antigen (Stefanová et al., 2002). It is unknown whether described differences between naïve CD4^+^ T cells of low and high self-reactivity are similarly dependent on subthreshold tonic self-pMHC signals. An alternative hypothesis is that naïve T cells are pre-wired by TCR signals obtained during thymic encounters with self-pMHC as they undergo positive selection, and that these differences are then epigenetically imprinted. Given our findings that some modulators of TCR signal strength remained distinct post activation, we examined which gene expression differences between CD5^lo^ and CD5^hi^ naïve CD4^+^ T cells were dependent on continuous self-pMHC interactions in the periphery, and which might be a result of thymic self-ligand interactions.

To investigate this, we sorted 15% CD5^lo^ and CD5^hi^ naïve CD4^+^ T cells and cultured them *ex vivo* in the absence of tonic self-pMHC interactions for 22 hours with IL-7 and performed bulk RNA-seq on these rested populations. After resting, sorted populations segregated into clusters distinct from their freshly isolated counterparts along PC1 (62% of the variation), while the differences between CD5^lo^ and CD5^hi^ naïve CD4^+^ T cells were preserved in PC2 (14% of the variation) (Figure 6A). As previously described, levels of CD5 itself, both at the protein- and transcript-level, rapidly decreased upon resting (Figure 6B) (Mandl et al., 2012; Smith et al., 2001). Interestingly, however, when sorted CD5^lo^ and CD5^hi^ naïve CD4^+^ T cells were rested, CD5 mRNA and protein expression remained distinct, despite decreasing from levels measured in freshly isolated cells (Figure 6C). Therefore, while CD5 expression levels are maintained by peripheral self-interactions, our data suggest that the retention of CD5 expression differences between CD5^lo^ and CD5^hi^ cells were independent of peripheral tonic self-ligand interactions.

**Figure 6.**
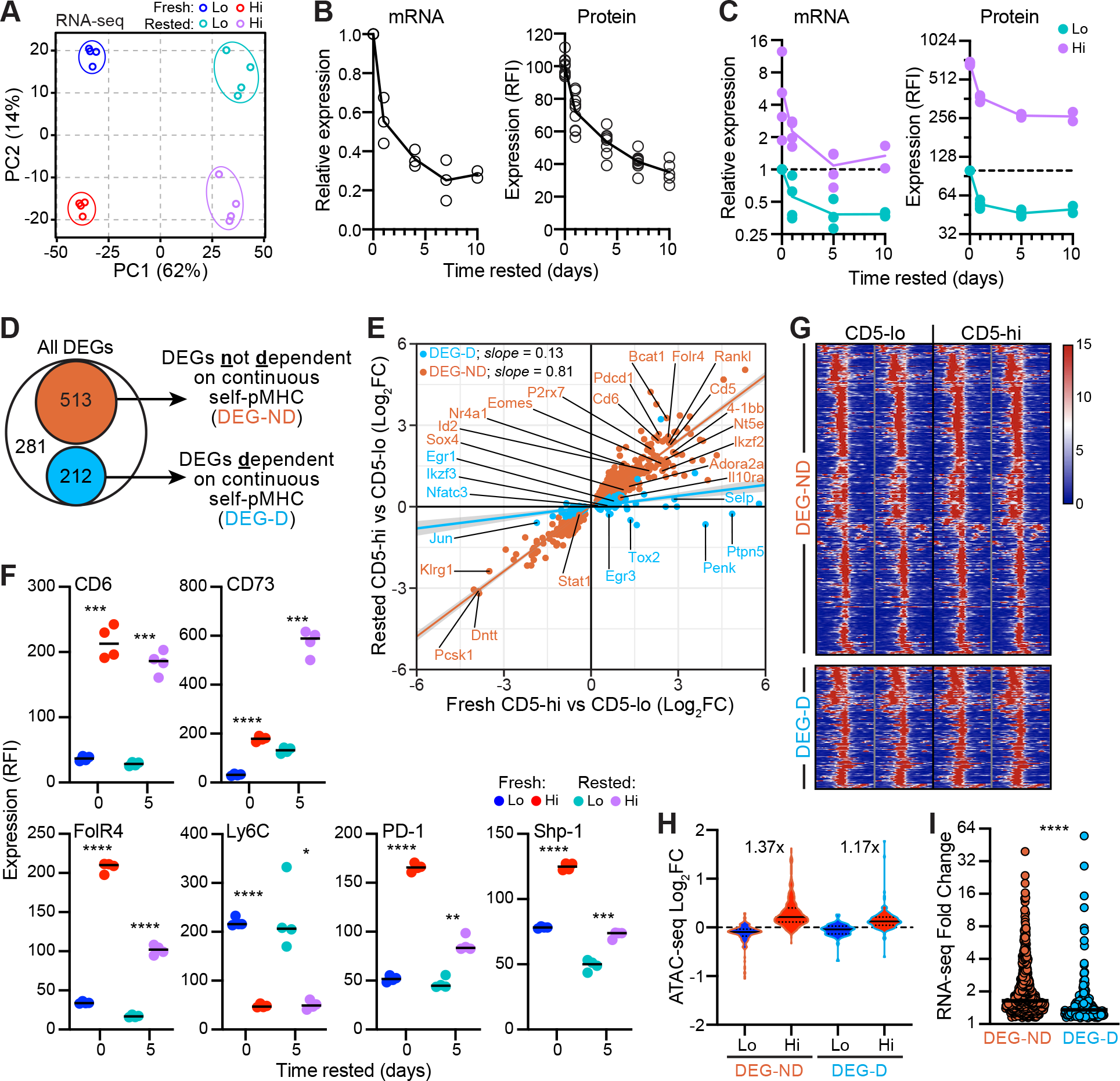
Withdrawal of naïve CD4^+^ T cells from self-pMHC identifies transcriptional and chromatin differences between CD5^lo^ and CD5^hi^ cells that do not rely on continuous self-pMHC interactions. (**A**) PCA of RNA-seq data from fresh and rested CD5^lo^ and CD5^hi^ naïve CD4^+^ T cells. (**B**) CD5 mRNA (normalized to *Gapdh*) and protein expression, relative to day 0, measured for naïve CD4^+^ T cells rested in dissociated culture in the presence of IL-7. (**C**) CD5 mRNA and protein expression determined after resting as in B of sorted 15% CD5^lo^ and CD5^hi^ naïve CD4^+^ T cells. Each data point represented 9-10 pooled mice; n=4 for Day 0 and 1; n=2 for Day 10. There was a significant effect between CD5^lo^ and CD5^hi^ in both mRNA (*) and protein (***) groups post-resting. (**D**) Venn diagram showing the division of CD5^hi^ versus CD5^lo^ DEGs identified by RNA-seq into two groups: DEG-ND (orange) were significantly differentially expressed (FDR*≤*0.01) in the fresh CD5^hi^ vs. CD5^lo^ bulk RNA-seq comparison and remained differentially expressed (FDR*≤*0.01) in the rested CD5^hi^ vs. CD5^lo^ RNA-seq comparison; DEG-D (blue) were significantly differentially expressed (FDR*≤*0.01) in the fresh CD5^hi^ vs. CD5^lo^ comparison but non-significant (FDR*≥*0.3) in the rested RNA-seq comparison. (**E**) FC expression of DEG-ND and DEG-D identified in D, in both the fresh and rested RNA-seq datasets. Lines indicate best fits to each DEG group, with slopes as shown. (**F**) Naïve CD4^+^ T cells were sorted into CD5^lo^ and CD5^hi^ populations and rested in dissociated culture in the presence of IL-7. RFI (normalized to total naïve CD4^+^ T cells) from genes identified from D. Data is summarized from 1 experiment. Each data point represents 3 mice pooled; lines denote group means. (**G**) ATAC-seq chromatin accessibility heatmap for open chromatin regions among DEG-ND and DEG-D. (**H**) Summary of data in G with fold differences between CD5^hi^ and CD5^hi^ open chromatin peaks indicated on the graph. (**I**) Gene expression fold changes from RNA-seq between CD5^hi^ and CD5^hi^ cells in DEG-ND and DEG-D groups identified in D. Lines denote group means. Statistics: Two-way ANOVA with Sidak’s multiple comparisons test (C), Paired t test (F), Mann-Whitney test (I). \**p*<0.05, \*\**p*<0.01, \*\*\**p*<0.001, \*\*\*\**p*<0.0001, ns = non-significant. See also Figure S6.

To examine whether other genes in our identified CD5^lo^ versus CD5^hi^ DEGs followed a similar pattern to *Cd5*, we compared expression levels in fresh compared to rested populations and designed classification criteria to subset genes into 2 groups: genes where expression differences between CD5-sorted populations were not dependent on continuous self-pMHC interactions (DEG-ND), or where differences were lost upon resting (i.e., expression differences were dependent on continuous self-pMHC interactions, DEG-D) (Figure 6D). A gene was classified as being within the DEG-ND subset if it was significantly differentially expressed at FDR≤0.01 in the fresh comparison and remained significant at FDR≤0.01 in the rested comparison. Conversely, to be classified as DEG-D, the gene had to be significantly differentially expressed at FDR≤0.01 in the fresh comparison but become non-significant at FDR≥0.3 in the rested comparison. With these criteria, we classified 513 of the 1006 DEGs as DEG-ND and 212 DEGs as DEG-D, while 281 DEGs did not fall into either group (Figure 6D). Plotting the CD5^lo^ versus CD5^hi^ fold-changes of the fresh versus the rested populations confirmed that, for the most part, DEG-ND genes fell on the line of equivalence, indicating that fold differences between CD5^lo^ versus CD5^hi^ naïve CD4^+^ T cells were unaffected by the absence of self-pMHC interactions, while DEG-D genes located approximately on a y=0 line, indicating that fold differences between CD5^lo^ versus CD5^hi^ converged to zero after resting (Figure 6E). These data suggested that transcriptional heterogeneity among the naïve CD4^+^ T cell population was explained both by differences which depended on continuous self-pMHC interactions and differences which were independent of tonic self-pMHC signals. Interestingly, most of the negative regulators of TCR signaling fell into the DEG-ND group (including *Bcat1*, *Nt5e*, *Adora2a*, *Pdcd1*, *Il10ra*, as well as *Cd6* and *Cd5*), consistent with their expression level being set during thymic development, possibly to prevent them from being negatively selected, as was the gene *Dntt* important in V(D)J recombination (Figure 6E). In addition, more than half of the T_FH_-associated genes found in our DEGs fell into the DEG-ND group, including *Icos*, *Itgb2*, *P2rx7*, and *Adora2a*, whereas 20% of TFH associated genes were found in the DEG-D group, including *Tox2* and *Dusp6*, consistent with the change in T_FH_ development we observed when CD4^+^ T cells experienced greater tonic TCR signal strength (**Figure S5A-D**). Of note, the TRs *Jun, Egr1, Egr3, Ikzf3*, and *Nfatc3* were all classified as DEG-D, while *Stat1, Eomes, Ikzf2, Id2, Sox4,* and *Nr4a1* were found in the DEG-ND group, indicating that even at the level of regulation of TRs some expression differences among naïve CD4^+^ T cells rely on continuous self-ligand interactions while others do not (Figure 6E). To verify whether the maintenance of differences among transcripts following resting was also observed at the protein level, we chose a subset involved in TCR signalling (CD6, CD73, FolR4, Ly6C, PD-1, and SHP-1) from the DEG-ND group and measured protein expression in sorted fresh and rested CD5^lo^ versus CD5^hi^ naïve CD4^+^ T cell populations (Figure 6F). To give protein levels time to turnover, we rested sorted cells for 5 days in dissociated culture and then compared expression of specific proteins in rested cells to baseline levels. We observed no loss in cell viability during this period (**Figure S6A**), and, as observed at the transcript level, while protein levels did change upon resting, the difference between CD5^lo^ and CD5^hi^ naïve CD4^+^ T cells was retained (Figure 6F**, Figure S6B**).

Our classification of genes into two distinct groups based on their reliance on continuous self-pMHC interactions suggested that at least a subset of the described transcriptionally-wired heterogeneity among naïve CD4^+^ T cells may reflect epigenetic differences established in the thymus. To test this hypothesis, we probed our ATAC-seq dataset to ask whether there were detectable differences in chromatin states between the genes in the two groups. We predicted that loci of DEG-ND genes would show a greater difference in accessible regions between CD5^lo^ and CD5^hi^ cells than would the loci of DEG-D genes. Indeed, on average, the fold difference in chromatin accessibility was greater in the DEG-ND compared to DEG-D group (Figure 6G **and 6H)**. In line with this, gene expression difference between CD5^lo^ and CD5^hi^ cells were greater in the DEG-ND set than the DEG-D set (Figure 6I). Collectively these data suggest that there are two sources of heterogeneity among naïve CD4^+^ T cells which impact their responsiveness to foreign antigen and their effector differentiation: differences that require continuous self-pMHC interactions in the periphery, and differences that do not require self-pMHC interactions, that may be epigenetically imprinted in the thymus.

## Discussion

CD4^+^ T cells play a critical role in orchestrating an immune response, with early fate decisions between T_FH_ and non-T_FH_ effector subset lineage decisions thought to be primarily determined by TCR engagement with cognate pMHC during priming (Ruterbusch et al., 2020). Here we showed that there are transcriptional and open-chromatin differences between CD4^+^ T cells that are present prior to their activation, are maintained after activation in the short-term, and impact TCR signal strength and early lineage choice upon antigen encounter. Further, our data imply that naïve CD4^+^ T cell heterogeneity is partly thymically imprinted and retained independent of interactions with self-pMHC in the periphery. Thus, our work reconciles prior studies that have described specific heterogeneous traits among naïve CD4^+^ T cells, not all of which were dependent on tonic TCR signals (Mandl et al., 2013; Matson et al., 2020; Persaud et al., 2014; Stefanová et al., 2002).

At the single cell level, our data highlighted that heterogeneity within the naïve CD4^+^ T cell pool is orchestrated by many interacting genes. We identified expression in modulators of TCR signaling as a key driver of heterogeneity among naïve CD4^+^ T cells, in addition to chromatin modifiers and genes involved in steady-state T cell trafficking between secondary lymphoid organs. Our RNA-seq data suggests that CD5 expression may be a better predictor of cellular behaviour at the population-level than at the single cell level. Indeed, population averages of sorted subsets of polyclonal naïve CD4^+^ T cells might therefore not always be mirrored when studying the behaviour of only a few specific TCR clonotypes and could explain some of the contradictory findings with regard to T_FH_-lineage differentiation biases described based on specific pairs of monoclonal TCR transgenic clones (Bartleson et al., 2020; Persaud et al., 2014).

Our findings extend on previous work that has implicated self-pMHC reactivity in the potential of CD4^+^ T cells to become IFN*γ*-producing T_H_1 cells, T_H_17 cells, or Tregs (Henderson et al., 2015; Martin et al., 2013; Sood et al., 2019), and suggests that the early lineage bifurcation into T_FH_ cells is also impacted by self-reactivity. Our data are consistent with recent work showing that T cells with longer dwell times between their TCR and pMHC on antigen presenting cells (and thus stronger TCR signaling) become IL-2 producing cells and signal in *trans* to IL-2-non-producing cells to reinforce non-T_FH_ effector differentiation (Ballesteros-Tato et al., 2012; DiToro et al., 2018). We found that increasing the strength of self-pMHC signals obtained by adoptive transfer of SMARTA TCR transgenic CD4^+^ T cells into T cell deficient mice led to a decrease in T_FH_ differentiation due to enhanced IL-2 production that is also modulated by TCR signal strength (Persaud et al., 2014). Thus, removing competitor T cells, by modulating both IL-2 and TCR signal strength, led to the opposite outcome with regard to T_FH_ differentiation predicted by TCR signal strength alone, and is in line with recent observations showing that Nur77^lo^ CD4^+^ T cells adoptively transferred into TCR*α*^-/-^ gave rise to a greater T_FH_ frequency than transferred Nur77^hi^ cells (Bartleson et al., 2020). Given recent data corroborating the use of CD5 as a marker for the self-ligand reactivity in human CD4^+^ T cells (Sood et al., 2021), it will be interesting to determine whether this relationship between self-reactivity and effector potential holds in humans.

While our data implicates thymically-imprinted epigenetic differences in the transcriptional heterogeneity among naïve CD4^+^ T cells, we did not address here whether such signals are impacted at different times during development. It is increasingly appreciated that T cell development is a layered process, with neonatally-derived T cells being distinct from adult-derived T cells in a number of ways (Adkins, 2003; Hebel et al., 2014; Mold et al., 2010; Rudd, 2020; Smith et al., 2018; Wang et al., 2010). Interestingly, in both mice and humans, CD5 expression on naïve CD4^+^ T cells is markedly lower in adults than in neonates (Dong et al., 2017; Lutes et al., 2021; Mandl et al., 2013). Moreover, CD4^+^ T cells derived from fetal or neonatal stem cells are more responsive to stimulation and are more likely to differentiate into Tregs and T_H_2 cells compared to adult stem cell derived CD4^+^ T cells (Adkins, 2003; Hebel et al., 2014; Mold et al., 2010; Wang et al., 2010). Thus, CD5^hi^ naïve CD4^+^ T cells could be enriched for fetal/neonatally-derived cells and it will be important to determine if this is a contributing factor to the diversity that we identified among naïve CD4^+^ T cells.

Whether differences among naïve CD4^+^ T cells can ultimately be related back to features of their specific TCRs remains an interesting and open question. It is intriguing that in CD5^lo^ naïve CD4^+^ T cells one of the top DEGs was *Dntt* (encoding TdT), as was also observed in other datasets of both mouse naïve CD8^+^ T cells and human naïve CD4^+^ T cells (Fulton et al., 2015; Sood et al., 2021). TdT is responsible for adding n-nucleotides during V(D)J recombination and thus diversifying the TCR repertoire (Benedict et al., 2000; Cabaniols et al., 2001). It has been proposed that CD5^hi^ T cells have a greater proportion of germline TCRs (lacking n-nucleotide insertions) with shorter complementarity-determining regions (CDR3s) and have been evolutionarily optimized to strongly bind to pMHC (Vrisekoop et al., 2014). It is possible that differences in *Dntt* expression play a role in dictating CDR3 length and self-pMHC reactivity. Ultimately, patterns in TCR sequences may exist that enable some prediction of self-pMHC reactivity and, therefore, the differentiation potential of individual T cell clones.

Together, our data shed light on which pre-existing transcriptome-level differences among naïve CD4^+^ T cells are accessible to interventions targeting self-pMHC peripheral interactions, compared to others that would require modulation at the chromatin level, which may aid in optimizing protocols for enhancing desirable T cell functions in clinical settings such as in adoptive cell therapies (Alspach et al., 2019; Borst et al., 2018).

## Acknowledgements

We would like to thank Geneviève Perreault, Patricia D’Arcy, and the animal facility staff at McGill for their excellent care of our animal colony. We are grateful to Ronald Germain (NIH), Troy Baldwin (University of Alberta), Nevil Singh (University of Maryland), Nienke Vrisekoop (Utrecht University), and members of the Mandl lab for their critical feedback and comments on earlier versions of the manuscript. We would also like to thank Camille Stegen and Julien Leconte at the McGill University Cell Vision Core Facility for performing all cell sorts. Computational analyses were enabled in part by support provided by Calcul Québec and Compute Canada (www.computecanada.ca). D.R. was funded by a Frederick Banting and Charles Best Canada Graduate Scholarships Doctoral Award (CIHR CGS-D) and a Tomlinson Doctoral Fellowship (McGill University). A.S. was funded by a Cole Foundation Postdoctoral Fellowship. D.L. and H.J.M. are junior 1 and junior 2 scholars of the FRQS, respectively. J.N.M. is a Canada Research Chair for Immune Cell Dynamics. This research was supported by a NSERC Discovery Grant (2016-03808) a McGill start-up fund to J.N.M., as well as a Cancer Research Society (22344) and a CIHR (PJT-168862) grant jointly to J.N.M and H.J.M. J.J.P.vB was funded by a NOW Vici grant (016.140.655).

## Author Contributions

D.R. and J.N.M. conceived and designed the project and research, with input from H.J.M. and J.T. D.R. performed most experiments with critical help from C.Sc., C.Sh., and D.W. Bulk RNA-seq was performed by J.N.M, with help from A.J.M. and J.S.T., and D.R. analyzed the bulk RNA-seq data. D.R., A.S. and M.L. generated the ATAC-seq dataset with input from H.J.M., and data was analysed by D.R. and H.W. The scRNA-seq data was generated and analysed by J.vB. and J.T. with input from D.R. and J.N.M. CD5 decay experiments were performed by P.A. The mutual information analysis was contributed by T.R. with input from P.F. Critical reagents and intellectual input were provided by S.A.C., M.J.R., L.B., P.F., and D.L. The manuscript was written by J.N.M. and D.R., with critical input from H.J.M and J.T., and feedback from all other authors.

## Materials and Methods

### Mice and infections

C57BL/6 mice, CD45.1^+^, Foxp3^GFP+^ transgenic (Oukka, 2007), SMARTA TCR transgenic (Oxenius et al., 1998), TCR*β*^-/-^ (Mombaerts et al., 1992), and MHCII^-/-^ mice (Madsen et al., 1999) were purchased from Jackson Laboratories (Bar Harbor, ME) and bred in-house. All mice were on a B6 background and used for experiments at 6-12 weeks of age. Animal housing, care and research were in accordance with the Guide for the Care and Use of Laboratory Animals and all procedures performed were approved by the McGill University, Maisonneuve-Rosemont Hospital Research Center, Radboud University, or NIAID Animal Care Committee. For *in vivo* infections, LCMV-Armstrong (LCMV) was propagated as previously described (Slifka and Whitton, 2001) and mice infected with 2×10^5^ plaque forming units (PFU) by intra-peritoneal injection (Wherry et al., 2003). Cellular responses were assessed 8 days post-infection.

### Lymphocyte isolation, resting, activation, and restimulation

Spleen and peripheral lymph nodes (inguinal, axillary, brachial, superficial cervical, and mesenteric) were isolated as previously described (Schneider et al., 2020). For experiments where naïve CD4^+^ T cells were rested in culture, cells (either total or sorted, as specified) were kept in complete RPMI (10% FBS, 1% L-glutamine, 1% pen/strep, 1% HEPES buffer, 1% Sodium Pyruvate, 1% non-essential Amino Acids, 0.1% 2-mercapto-ethanol 1000X solution) supplemented with IL-7 (10 ng/mL, Biolegend). To activate T cells, sorted cells or total splenocytes were cultured in complete RPMI in flat-bottomed 96-well plates coated with *α*-CD3 and *α*-CD28 (Invitrogen; both at 3μg/mL). Restimulation of splenocytes for cytokine production was performed as previously described (Schneider et al., 2020).

### Flow cytometry

Samples were incubated in Fixable Viability Dye (AF780 or eF506, eBioscience) diluted in PBS for 20 minutes at 4°C. Extracellular antibodies were diluted in FACS buffer (2% FBS and 5mM EDTA in PBS) with Fc Block (eBioscience) and incubated for 30 minutes at 4°C. Samples requiring intracellular staining were subsequently incubated in FoxP3 Transcription Factor Fixation/Permeabilization Concentrate and Diluent (Life Technologies) for 30 minutes at 4°C. Intracellular antibodies were diluted in permeabilization wash buffer and incubated for 30-60 minutes at 4°C. Directly conjugated antibodies used were as follows: TCR*β* (H57-597), CD4 (RM4.5), CD8 (53-6.7), CD5 (53-7.3), Foxp3 (FJK-16s), CD44 (IM7), CD62L (MEL-14), CD25 (PC61.5), CD45 (30F11), CD98 (RL388), Gitr (DTA-1), LFA-1 (H155-78), CD73 (TY/11.8), PD-1 (29F.1A12), FolR4 (eBio12A5), Ly6C (HK1.4), CD6 (OX-129), CXCR5 (SPRCL5), CD45.1 (A20), CD45.2 (104), TOX (TXRX10), CD69 (H1.2F3), B220 (RA3-6B2), F4/80 (BM8), Ly6G (1A8), CD11b (M1/70), CD11c (N418), NK1.1 (PK136), CD19 (eBio1D3), and IL-2 (JES6-5H4). Primary unconjugated antibodies used were: LEF-1 (C12A5), TCF-1 (C46C7), SHP-1 (C14H6). Secondary conjugated antibodies used were either Goat anti-Rabbit IgG (H+L) Alexa Fluor 488 or Donkey anti-Rabbit IgG (H+L) Alexa Fluor 647. For samples assessed for SHP-1 expression, cells were fixed with 1X TFP Fix/Perm Buffer for 50 minutes at 4°C, then incubated in Perm Buffer III (BD Biosciences) for 20 minutes on ice. Fc Block, surface, and intracellular antibodies were diluted in 1X TFP Perm/Wash Buffer and incubated for 50 minutes at 4°C, and secondary antibody diluted in 1X TFP Perm/Wash Buffer was added for an additional 50 minutes at 4°C. For all flow cytometry experiments, cells were acquired using an LSRFortessa (BD Bioscience) and analyzed with FlowJo software (BD Bioscience).

### Cell sorts

Lymphocytes from B6 or CD45.1^+^ congenic mice were isolated in single cell suspension as described. Samples for bulk RNA-seq, ATAC-seq, *in vivo* or *in vitro* assays were pooled from spleen and lymph nodes (inguinal, axillary, brachial, mesenteric, and cervical) from 2-10 mice. Samples for scRNA-seq were from a spleen from a single mouse. Total isolated cells or cells magnetically enriched for CD4 or total T cells (Stemcell EasySep mouse total T cell or CD4^+^ T cell enrichment kits) were then incubated in fixable viability dye, and subsequently stained with surface antibodies for 1 hour at 4°C. Naïve CD4^+^ T cell were sorted on CD5 expression (top and bottom 15%) for bulk analyses; single naïve CD4^+^ T cells were sorted into 384-well plates for subsequent scRNA-seq. Naïve CD4^+^ T cells were sorted on singlets, live, dump-negative (RNA-seq and ATAC-seq), TCR*β*^+^ (bulk- and scRNA-seq), CD4^+^, CD8^-^, CD25^-^ (scRNA-seq and *in vivo* and in *vitro* assays) or Foxp3^GFP+^ (RNA-seq), CD44^lo^, CD62L^hi^, and 15% CD5^lo^ and CD5^hi^ (RNA-seq, ATAC-seq, and *in vivo* and *in vitro* assays). Dump channel included B220, CD11b, CD11c, F4/80, Ly6G, NK1.1, and CD69 for RNA-seq; the ATAC-seq dump channel also included CD19 and CD25 (ATAC-seq). Sorts were performed on either a FACS Aria Fusion, Aria III, or Aria II SORP (BD Bioscience). All cell populations were sorted to >90% purity for bulk populations.

### Adoptive cell transfers

For all adoptive cell transfer experiments, donors and recipients were sex-matched.

#### LCMV infection

15-18 CD45.1^+^ or CD45.2^+^ mice were used as donors to obtain a total of 12-20×10^6^ cells from each of 15% CD5^lo^ and 15% CD5^hi^ cells sorted as detailed above. 6-10×10^6^ sorted donor cells were adoptively transferred by i.v. injection into CD45.2^+^ or CD45.1^+^ recipients (n=2 per group), respectively. One day post-transfer, mice were infected with LCMV as described. Cells were isolated from the spleens and peripheral lymph nodes of recipient mice 8 days post-infection.

#### SMARTA transgenic T cell adoptive transfer

1×10^4^ CD45.2^+^ SMARTA CD4^+^ T cells were adoptively transferred by i.v. injection into CD45.1^+^ recipients. One day post-transfer, mice were infected with LCMV as described. Cells were isolated form the spleens of recipient mice 8 days post-infection.

### Bulk RNA sequencing

1×10^6^ cells from four independent samples, each with cells pooled from 2 mice, were sorted as described and CD5^lo^ and CD5^hi^ naïve CD4^+^ T cells were either directly added to 500μL TRIzol (ThermoFisher Scientific) or rested in complete RPMI supplemented with IL-7 for 22 hrs first. RNA was purified using RNA miniprep kit (Zymo Research) according to manufacturers’ recommendations. 500ng of purified RNA was used to prepare RNA-seq libraries using TruSeq mRNA library preparation kit v2 (Illumina). Libraries were sequenced on an Illumina HiSeq 2000 using v3 chemistry and 50 cycle paired end reads. Illumina bcl files were converted to FASTQ using CASAVA1.8.2 and mapped to the UCSC mm9 mus musculus genome annotation using Tophat 2.0.11 (Kim et al., 2013). Reads overlapping exons were counted using featureCounts version 1.4.5 from the SubRead package (Liao et al., 2014), with a minimum read mapping quality score of 10. Normalized read counts differential gene expression analysis was performed with EdgeR (Robinson et al., 2010). In order for a gene to be included in the matrix a minimum CPM value of 5 in at least 3 of the 4 replicates was required. The p-values were corrected using the Benjamini-Hochberg method and a FDR threshold of <0.01 was considered significant. No fold change threshold was set unless stated otherwise.

### Single cell RNA sequencing

Each well within a 384-well plate contained CEL-Seq2 primers covered by mineral oil. Primers consisted of a 24bp polyT stretch, a 6bp random molecular barcode (UMI), a cell-specific barcode, the 5’ Illumina TruSeq small RNA kit adaptor and a T7 promoter. After sorting, the plates were frozen at −80°C until further use. Single cell RNA-seq library preparation and sequencing was performed by Single Cell Discoveries (Utrecht, Netherlands) (Artegiani et al., 2017). Libraries were prepared following the SORT-seq protocol (Muraro et al., 2016), which consists of an automated and improved version of the CEL-Seq2 protocol (Hashimshony et al., 2016). Briefly, cells were first lysed for 5 minutes at 65°C, and reverse transcription and second strand mixes were dispensed by the Nanodrop II liquid handling platform (GC Biotech). Single cell double stranded cDNAs were pooled together and *in vitro* transcribed for linear amplification. Illumina sequencing libraries were prepared using the TruSeq small RNA primers (Illumina) and sequenced paired-end at 75 bp read length the Illumina NextSeq. Paired-end reads from Illumina sequencing were aligned to the mouse transcriptome genome by BWA (Li and Durbin, 2010). Read 1 contained the barcode information and was used for assigning reads to correct cells and libraries, while read 2 was mapped to gene models. Reads that mapped equally well to multiple locations were discarded. Read counts were first corrected for UMI barcode by removing duplicate reads that had identical combinations of library, cell-specific, and molecular barcodes and were mapped to the same gene. For each cell barcode the number of UMIs for every transcript was counted, and transcript counts were then adjusted to the expected number of molecules based on counts, 256 possible UMI’s and poissonian counting statistics (Grün et al., 2014). A unique feature of this protocol is the combination of both flow cytometry staining and RNA sequencing; this allowed for the simultaneous tracking of select protein expression and gene expression on single cells.

### Single-cell RNA-seq data analysis

Raw read counts were first subjected to quality control. We identified two blocks of wells with fewer than 500 non-spike-in reads. To exclude these and similar low-content wells, we applied a UMAP clustering on all wells (including the spike-in reads) and excluded the cluster of cells that was mainly composed by the low-read wells. After quality control, 697 wells out of 1152 were kept in the analysis. R packages ‘scater’ (McCarthy et al., 2017) and ‘scran’ (Lun et al., 2016) were used for further processing, the spike-in reads were removed and expression values were normalized to library size and normalized log expression values and gene variance were determined as described previously described (Lun et al., 2016). Mitochondrial genes were excluded before modeling gene variance. The processed data was then plotted or subjected to UMAP clustering (McInnes et al., 2020), performed with the R package ‘uwot’ (Melville, 2019) using the cosine distance and a neighbourhood size of 30. For defining CD5 low, mid, and high cells the CD5 mean fluorescent intensity was logged by the flow cytometer when we sorted the cells into individual wells.

### ATACseq library preparation, sequencing, and visualization

Two independent biological replicates of CD5^lo^ and CD5^hi^ naïve CD4^+^ T cells were sorted as described, counted and 1×10^5^ nuclei pelleted. ATAC-seq libraries were prepared from the fresh nuclei pellets by the Institut de recherches cliniques de Montréal. Briefly, paired-end 42bp sequencing reads were generated by Illumina sequencing (using a NovaSeq6000). The quality of the sequenced reads was checked using FastQC tool v0.10.1 (Babraham Bioinformatics), and low-quality bases removed using Trimmomatic v.0.33 (Bolger et al., 2014). The trimmed reads were mapped to the mouse UCSC mm9 genome using Bowtie 1.0.0 (Langmead et al., 2009), in paired-end mode with --best parameter. Peak calling was performed using MACS1.4.1 (Zhang et al., 2008) with p-values <10^-7^. Subpeaks were identified using PeakAnalyzer (Salmon-Divon et al., 2010), with parameters: *valley*=0.5 and *cutoff*=5 counts per million (cpm). Normalized sequenced read density profiles (bigwig) were generated using makeUCSCfile from Homer package (Heinz et al., 2010), normalizing the total number of reads in each sample to 10^6^, and visualized on Integrative Genomics Viewer (IGV) (Thorvaldsdóttir et al., 2013). Peaks identified in the biological replicates were pooled using mergePeaks from Homer package, merging peak summits within 50bp to each other. Read densities around the peak summits were retrieved using annotatePeaks from Homer package and quantiles normalized for fold change comparison between CD5^lo^ and CD5^hi^ replicates (Bolstad, 2020). Transcription factor binding motif enrichment analysis was performed using Homer package on unique peaks found only in CD5^lo^ or CD5^hi^ replicates with a *P*-value <10^-4^. Hierarchical clustering of the peaks near the DEG-ND and DEG-D gene sets were performed using Pearson correlation with complete linkage method.

### Analyses and Statistics

#### Heatmaps

For RNA-seq these were created by either showing individual replicates or average expression within replicates. Log_10_(CPM+1) were visualized using the pheatmap package in R-Project on a color scale of black-blue-white-orange-red (Raivo, 2018). For ATAC-seq heatmaps were created using annotatePeaks from Homer package, taking read densities ±1kb with bin size of 50bp for the highest peak summit near each gene TSS. Images were generated using MeV tool with blue-white-red scale (Howe et al., 2011).

#### Geneset enrichment analysis (GSEA)

GSEAs were performed as previously described (Subramanian et al., 2005) using gene sets defined by the Molecular Signatures Database (Liberzon et al., 2015) or otherwise described.

#### Gene ontology pathway analysis

Enrichment of GO terms in naïve CD4^+^ T cells was performed using ClueGO (verson 2.5.4) (Bindea et al., 2009). The following parameters were used when running ClueGO on the top 5% most variable genes from the scRNA-seq: Min GO Level =4, Max GO Level =6, Minimum Number of Genes associated to GO term =6, and Minimum Percentage of Genes associated to GO term =6. The following parameters were used when running ClueGO on bulk RNA-seq DEGs: Min GO Level =3; Max GO Level =4. For CD5^lo^ cells: Minimum Number of Genes associated to GO term =3; Minimum Percentage of Genes associated to GO term =5. For CD5^hi^ cells: Minimum Number of Genes associated to GO term =20; Minimum Percentage of Genes associated to GO term =10. Enrichment p-values were based on a hypergeometric test and Benjamini-Hochberg method used for multiple testing correction. For bulk RNA-seq only pathways with *P ≤* 0.05 were considered significant.

#### PCA

PCA plots were built using filtered log_2_CPM (RNA-seq) or log_2_-transformed read densities around peak summits (ATAC-seq) using ggplot2 package in R-Project (Wickham, 2016).

#### Mutual Information

Mutual Information (MI) is a robust, non-parametric measure of the statistical relationship between observables with distinct advantages over simple correlation measures (Chan et al., 2017). MI is computed as:

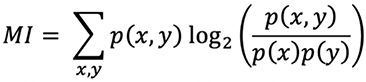

(Joint) probability distributions are obtained by binning the data into 96 geometrically spaced bins over the full Mean Fluorescent Intensity (MFI) range (10^0^ – 10^6^) for TCR*β*, CD5, CD6, and SHP-1.

#### ScatterSlice analysis

Scale values corresponding to single CD4^+^ T cells with expression of TCR*β*, Ly6C, CD5, CD6, TOX, and SHP-1 from flow cytometry data were identified and exported as csv files for analysis in the R-Project ScatterSlice (Cotari et al., 2013). Cells were divided into defined bins (15×15 matrix with a minimum of 15 cells per bin) and within each bin, the average MFI of Shp-1, Ly6C, TOX, or CD6 was projected in false-color onto a plot of TCR*β* versus CD5 expression.

#### Statistical analyses

Group comparisons were performed using Prism V9 (GraphPad). Unless specified, data are presented as mean ± standard deviations (SD) with each data point representing an individual mouse. The cut-off for significance considered was *p*≤0.05 for all analyses unless otherwise stated.

## Resource Availability

### Lead Contact

Further information and requests for resources and reagents should be directed to and will be fulfilled by the lead contact, Judith Mandl.

### Materials Availability

This study did not generate new unique reagents.

### Data and Code Availability

The data reported in this paper have been deposited in the Gene Expression Omnibus (GEO) database under accession numbers GSEXXXXX (bulk RNA-seq), GSEXXXXX (bulk ATAC-seq) and GSEXXXXX (scRNA-seq). Full source code for all quality controls and analysis steps for the scRNA-seq is available via public GitHub repository at https://github.com/jtextor/cd5-scrna.

## Supplemental Information

**Table S1.** Counts matrix for transcripts in scRNA-seq analysis performed on individual naive CD4^+^ T cells. CD5 surface protein expression (cd5.level) was measured by flow cytometry for each cell. Only cells that passed quality control are included.

**Table S2.** List of top 5% most variable genes among naïve CD4^+^ T cells from single cell RNA-seq. Log2 normalized counts, gene expression variance, and gene classification are indicated.

**Table S3.** List of DEGs identified in comparing CD5^lo^ and CD5^hi^ naïve CD4^+^ T cells (FDR < 0.01) using bulk RNA-seq. Log2 fold changes, absolute fold changes, log2 CPMs and p-values are indicated. Gene group classifications defined in Figure 6 are indicated in column G.

**Table S4.** Counts matrix for all detected transcripts in bulk RNA-seq in sorted fresh and rested CD5^lo^ and CD5^hi^ naïve CD4^+^ T cells.

