## Supplemental Figures for "Pre-existing chromatin accessibility and gene expression differences among naïve CD4^+^ T cells influence effector potential"

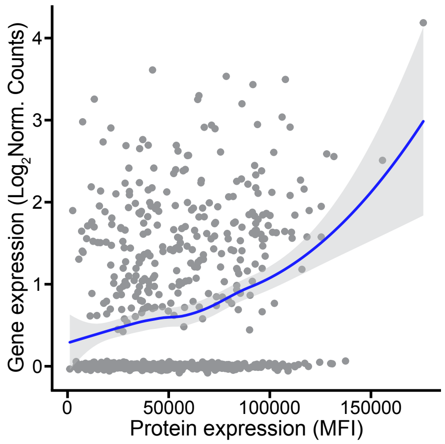


**Figure S1. CD5 expression among single naïve CD4$^+$ T cells. Related to Figure 1.** Comparison of *Cd5* gene expression from scRNA-seq (y-axis) and CD5 protein expression from corresponding sort (x-axis) for naïve CD4$^+$ T cells. Each data point represents a single cell.

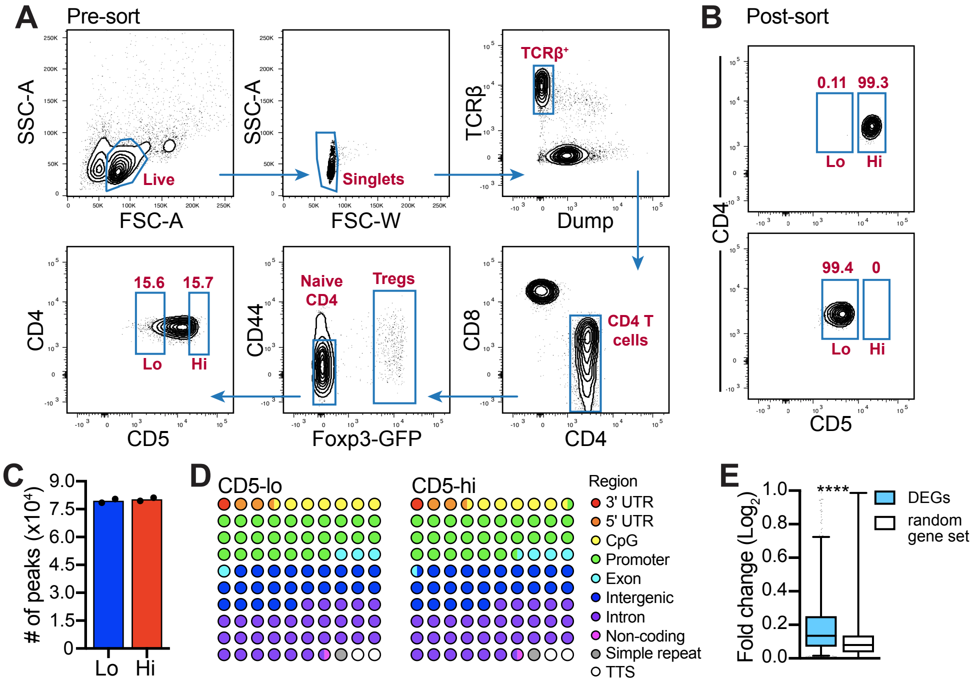


**Figure S2. Sort strategy and differences in chromatin accessible regions between CD5$^{lo}$ and CD5$^{hi}$ naïve CD4$^{+}$ T cells. Related to Figure 2.** (**A**) Flow cytometry plots illustrating the sort strategy used for RNA- and ATAC-seq. Dump channel included B220, CD11b, CD11c, F4/80, Ly6G, NK1.1, and CD69. (**B**) Representative post-sort purity for RNA- and ATAC-seq; numbers above gates indicate percent of cells in gate. (**C-D**) Global analysis of DARs in sorted CD5$^{lo}$ and CD5$^{hi}$ naïve CD4$^{+}$ T cells from ATAC-seq. Total number of peaks (C), proportion of peaks annotated to genomic regions (D). (**E**) Absolute fold expression of DARs from identified DEGs between CD5$^{hi}$ versus CD5$^{lo}$ as compared DARs corresponding to a random set of genes. Box plot shows the quartiles and the 5th and 95th percentiles as whiskers. Statistics: Mann-Whitney test (E).****$p$<0.0001.

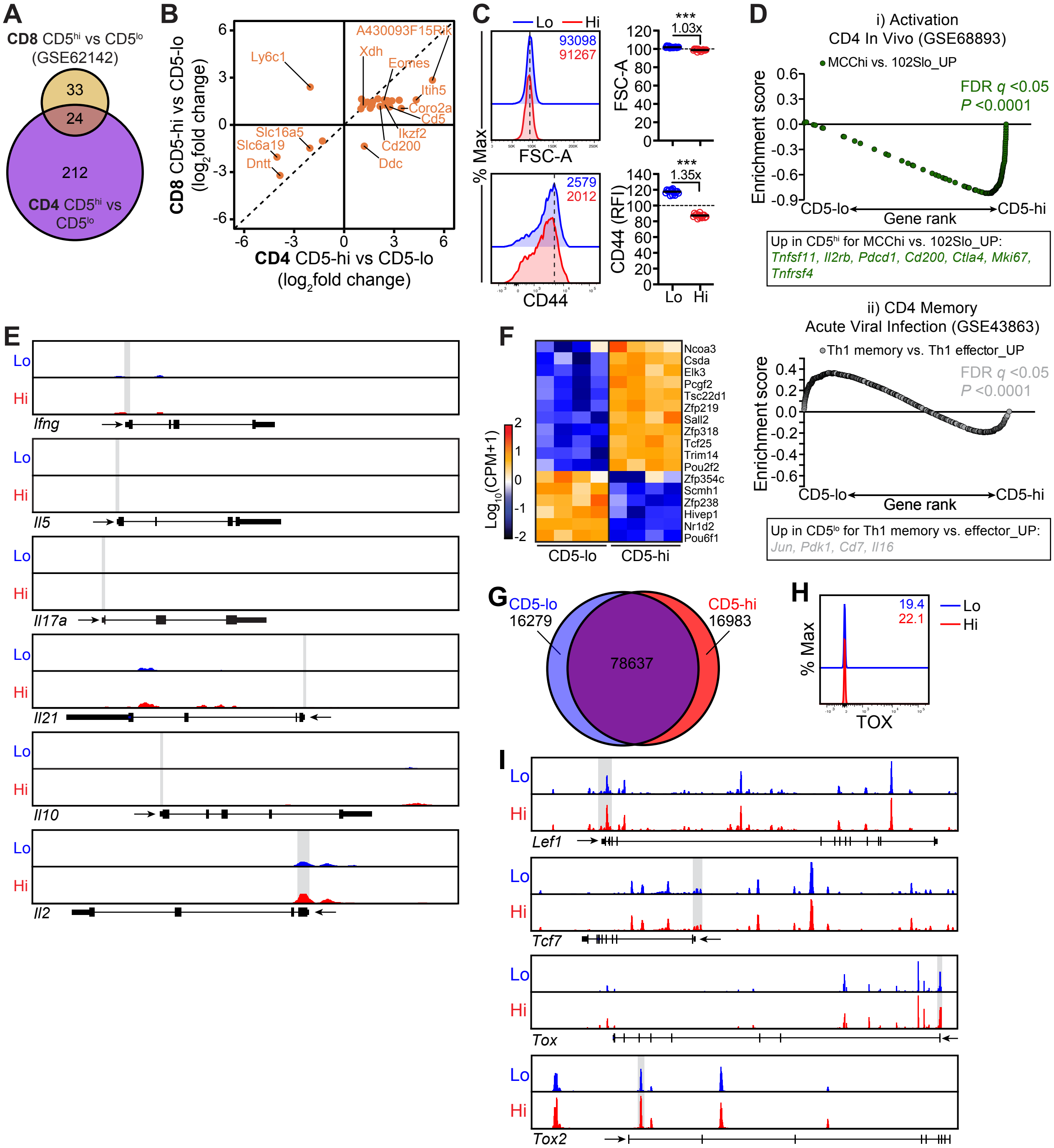

A
CD8 CD5hi vs CD5lo
(GSE62142)
33
24
212
CD4 CD5hi vs CD5lo
B
CD8 CD5-hi vs CD5-lo (log2fold change)
CD4 CD5-hi vs CD5-lo (log2fold change)
A430093F15Rik
Ly6c1
Xdh
Eomes
Itih5
Coro2a
Cd5
Ikzf2
Cd200
Ddc
Slc16a5
Slc6a19
Dntt
C
Lo
Hi
93098
91267
FSC-A
% Max
2579
2012
CD44
***
1.03x
FSC-A
***
1.35x
CD44 (RFI)
Lo
Hi
D
i) Activation
CD4 In Vivo (GSE68893)
MCChi vs. 102Slo_UP
FDR q <0.05
P <0.0001
Enrichment score
CD5-lo
Gene rank
CD5-hi
Up in CD5hi for MCChi vs. 102Slo_UP:
Tnfsf11, Il2rb, Pdcd1, Cd200, Ctla4, Mki67, Tnfrsf4
ii) CD4 Memory
Acute Viral Infection (GSE43863)
Th1 memory vs. Th1 effector_UP
FDR q <0.05
P <0.0001
Up in CD5lo for Th1 memory vs. effector_UP:
Jun, Pdk1, Cd7, Il16
E
Lo
Hi
Ifng
Il5
Il17a
Il21
Il10
Il2
F
Log10(CPM+1)
Ncoa3
Csda
Elk3
Pcgf2
Tsc22d1
Zfp219
Sall2
Zfp318
Tcf25
Trim14
Pou2f2
Zfp354c
Scmh1
Zfp238
Hivep1
Nr1d2
Pou6f1
CD5-lo
CD5-hi
G
CD5-lo
16279
CD5-hi
16983
78637
H
19.4
22.1
% Max
TOX
I
Lef1
Tcf7
Tox
Tox2

**Figure S3. Comparison of chromatin accessible regions and transcriptomes of CD5$^{hi}$ versus CD5$^{lo}$ CD4$^{+}$ T cells. Related to Figure 3.** (**A**) Venn diagram of DEGs identified in **Fig. 2** with FC≥2, compared to published DEGs determined for naïve CD8$^{+}$ T cells sorted into CD5$^{hi}$ and CD5$^{lo}$ (GSE62142). (**B**) Fold change expression differences between CD5$^{hi}$ and CD5$^{lo}$ of the 24 DEGs identified in the venn diagram overlap in A. (**C**) FSC-A and CD44 protein expression levels of CD5$^{lo}$ and CD5$^{hi}$ naïve CD4$^{+}$ T cells (CD44 RFI is relative to total naïve CD4$^{+}$ T cells). Representative flow cytometry histograms are shown and data summarized from 3 independent experiments. Dotted lines in histograms denote CD5$^{lo}$ modes; data point in graphs represent individual mice; lines denote group means. (**D**) GSEA showing enrichment of gene signatures in (i) CD4$^{+}$ T cell activation *in vivo*, GSE68893, and (ii) CD4$^{+}$ T cell memory during acute LCMV infection, GSE43863, relative to the CD5$^{lo}$ versus CD5$^{hi}$ naïve CD4$^{+}$ T cell comparison. (**E**) ATAC-seq signal profiles of T cell effector cytokines from one of 2 independent experiments. Promoter regions are highlighted in grey. (**F**) Heatmap of differentially expressed transcription factors (TF) between CD5$^{lo}$ and CD5$^{hi}$ naïve CD4$^{+}$ T cells not classified in **Fig. 3D**. (**G**) Venn diagram of unique DARs from ATAC-seq for CD5$^{lo}$ and CD5$^{hi}$ naïve CD4$^{+}$ T cells used for TF binding motif enrichment analysis in **Fig. 3E**. (**H**) Representative flow cytometry histogram of TOX FMO control in gated CD5$^{lo}$ and CD5$^{hi}$ naïve CD4$^{+}$ T cells. (**I**) ATAC-seq signal profiles of *Lef1*, *Tcf7*, *Tox*, and *Tox2* gene loci from one of 2 independent experiments. Promoter regions are highlighted in grey. Statistics: Wilcoxon matched-pairs signed rank test (C). ***$p$<0.001.

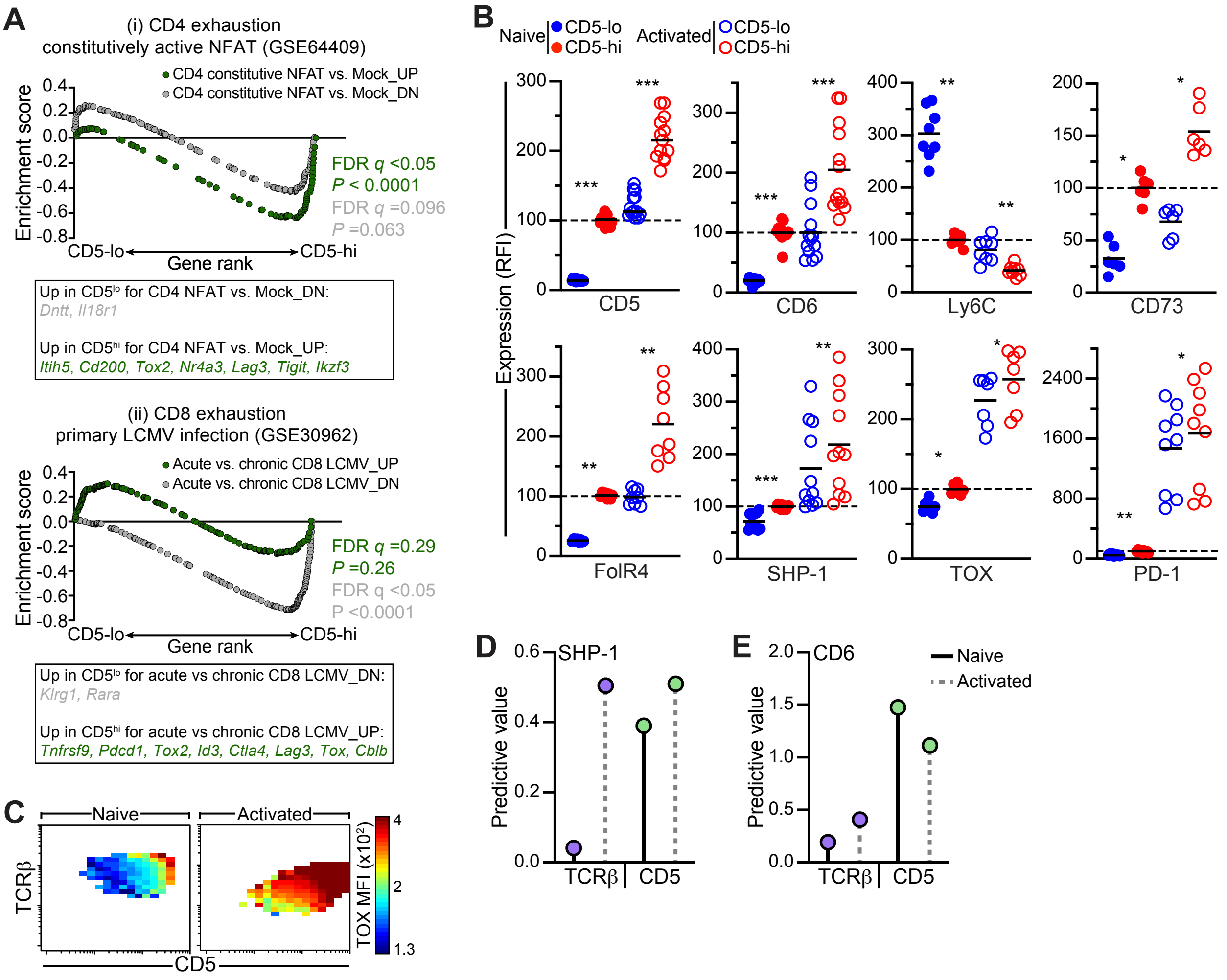


**Figure S4. CD5 expression on naïve CD4+ T cells predicts divergence of functional differences post-activation. Related to Figure 4.** (**A**) GSEA of gene signatures in mutant/constitutively-active form of NFAT1 overexpressing CD8$^{+}$ T cells (GSE64409) and T cell exhaustion during chronic viral infection (GSE30962) in CD5$^{lo}$ or CD5$^{hi}$ naïve CD4$^{+}$ T cells. (**B**) Protein expression (relative to sorted CD5$^{hi}$ naïve CD4$^{+}$ T cells) measured by flow cytometry in sorted 15% CD5$^{lo}$ and CD5$^{hi}$ naïve CD4$^{+}$ T cells pre- and 24 hours post-activation with anti-CD3/CD28. Activated cells were gated on CD44$^{hi}$CD25$^{+}$ or CD44$^{hi}$CD62L$^{-}$. Data summarized from 2-4 independent experiments; individual data points represent 2-5 pooled mice; lines denote group means. (**C**) 3-dimensional single-cell flow cytometry analysis for TOX expression among naïve CD4$^{+}$ T cells pre- and 24 hours post-activation. Color scale represents MFI of TOX in bins of at least 10 cells across the full protein expression spectrum of CD5 (x-axis) and TCRβ (y-axis). (**D,E**) Mutual information analysis to determine the importance of CD5 or TCRβ in predicting SHP-1 (D) or CD6 (E) expression levels in naïve CD4$^{+}$ T cells pre- and 24 hours post-activation. Statistics: Wilcoxon matched-pairs signed rank test (B). *$p$<0.05, **$p$<0.01, ***$p$<0.001.

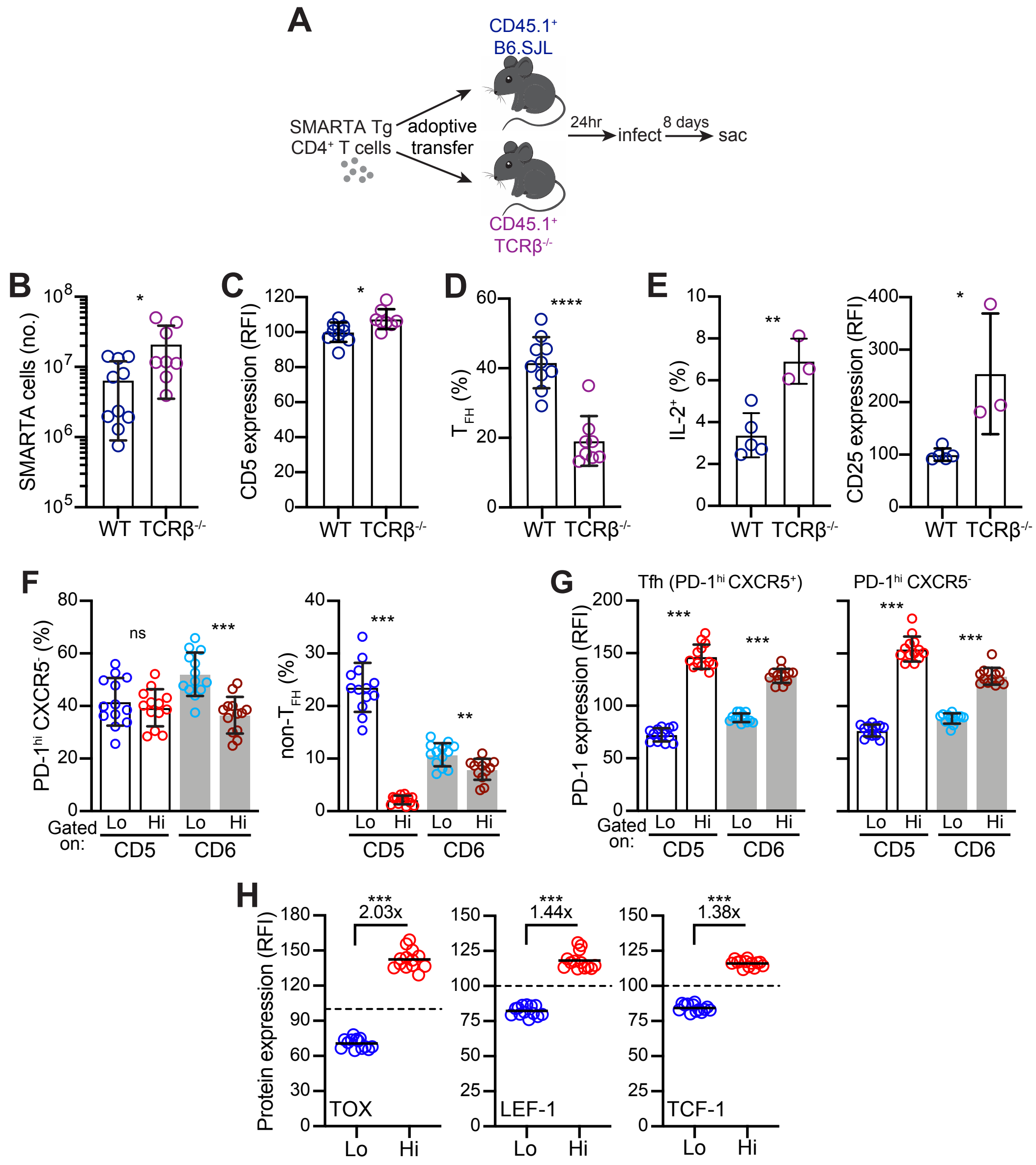


**Figure S5. High CD5 and CD6 expression identifies CD4+ T cells which develop into $T_{FH}$ cells with higher PD-1 and TOX expression. Related to Figure 5.** (**A-E**) Congenically labelled (CD45.2+) SMARTA transgenic CD4+ T cells were tranferred into WT or TCRβ-/- mice that were infected 1 day later with LCMV (A). Total number of activated (CD44hi) transferred SMARTA cells (B). CD5 protein expression (relative to SMARTA cells transferred into WT mice) of transferred SMARTA cells (C). Percent $T_{FH}$ of transferred SMARTA cells (D). Percent IL-2+ and CD25 protein expression (relative to SMARTA cells transferred into WT mice) of transferred SMARTA cells (E). All data are summarized from 1-2 independent experiments; each data point is from an individual mouse. (**F**) Activated (CD44hi) CD4+ T cells isolated on day 8 post LCMV infection were gated on the top and bottom 15% CD5- or CD6-expressing cells and the percent PD-1hiCxcr5- and non-$T_{FH}$ (PD-1loCxcr5-) determined. (**G**) PD-1 RFI (relative to total $T_{FH}$ cells) of gated cells from F. (**H**) TOX, LEF-1, and TCF-1 protein expression (relative to total $T_{FH}$ cells) in $T_{FH}$ cells from gated CD5lo and CD5hi CD4+ T cells. Statistics: Unpaired t test (B-E) and Wilcoxon matched-pairs signed rank test (F-H). **p*<0.05, ***p*<0.01, ****p*<0.001, *****p*<0.0001, ns= non-significant.

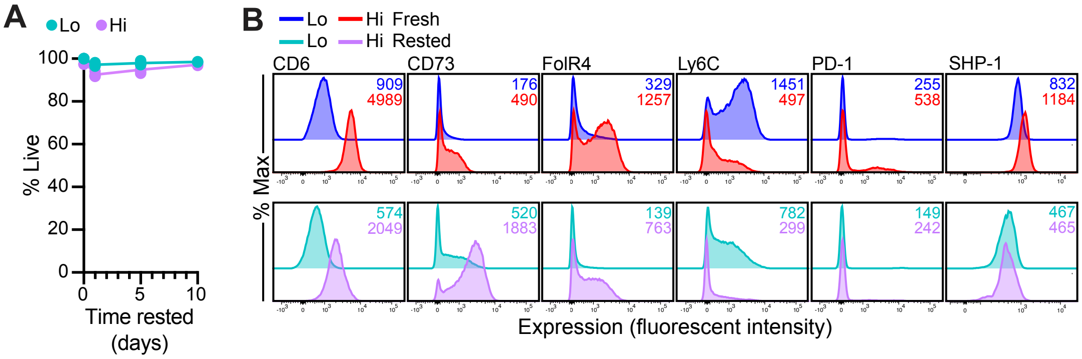


**Figure S6. Withdrawal of naïve CD4$^+$ T cells from self-pMHC identifies proteins that do not rely on continuous self-pMHC interactions. Related to Figure 6.** (**A**) Naïve CD4$^+$ T cells were sorted into 15% CD5$^{lo}$ and CD5$^{hi}$ populations, rested in dissociated culture in the presence of IL-7 for up to 5 days, and cell viability determined. Data is summarized from 1 experiment; each data point is from an individual mouse, n=4 for Day 0, 1, and 5; n=2 for Day 10. (**B**) Representative flow cytometry plots of proteins from sorted CD5$^{lo}$ and CD5$^{hi}$ naïve CD4$^+$ T cells rested in dissociated culture in the presence of IL-7 for 5 days. Numbers in top right of histograms indicate fluorescence intensity in fresh or rested CD5$^{lo}$ and CD5$^{hi}$ populations. Statistics: Two-way ANOVA with Sidak's multiple comparisons showed a significant effect ($p$<0.01) between CD5$^{lo}$ and CD5$^{hi}$ naive CD4 T cells at Day 1, but not at Day 0, 5, or 10 (A) .
